# Single-cell multi-region dissection of brain vasculature in Alzheimer’s Disease

**DOI:** 10.1101/2022.02.09.479797

**Authors:** Na Sun, Leyla Anne Akay, Mitchell H. Murdock, Yongjin Park, Adele Bubnys, Kyriaki Galani, Hansruedi Mathys, Xueqiao Jiang, Ayesha P. Ng, David A. Bennett, Li-Huei Tsai, Manolis Kellis

**Author notes:** These authors jointly supervised this work: Manolis Kellis, Li-Huei Tsai.

## Abstract

Cerebrovascular breakdown occurs early in Alzheimer’s Disease (AD), but its cell-type-specific molecular basis remains uncharacterized. Here, we characterize single-cell transcriptomic differences in human cerebrovasculature across 220 AD and 208 control individuals and across 6 brain regions. We annotate 22,514 cerebrovascular cells in 11 subtypes of endothelial, pericyte, smooth muscle, perivascular fibroblast, and ependymal cells, and how they differ in abundance and gene expression between brain regions. We identify 2,676 AD-differential genes, including lower expression of PDGFRB in pericytes, and ABCB1 and ATP10A in endothelial cells. These AD-differential genes reveal common upstream regulators, including MECOM, EP300, and KLF4, whose targeting may help restore vasculature function. We find coordinated vasculature-glial-neuronal co-expressed gene modules supported by ligand-receptor pairs, involved in axon growth/degeneration and neurogenesis, suggesting mechanistic mediators of neurovascular unit dysregulation in AD. Integration with AD genetics reveals 125 AD-differential genes directly linked to AD-associated genetic variants (through vasculature-specific eQTLs, Hi-C, and correlation-based evidence), 559 targeted by AD-associated regulators, and 661 targeted by AD-associated ligand-receptor signaling. Lastly, we show that APOE4-genotype associated differences are significantly enriched among AD-associated genes in capillary and venule endothelial cells, and subsets of pericytes and fibroblasts, which underlie the vascular dysregulation in APOE4-associated cognitive decline. Overall, our multi-region molecular atlas of differential human cerebrovasculature genes and pathways in AD can help guide early-stage AD therapeutics.

## Introduction

The blood-brain-barrier (BBB) separates brain parenchyma from peripheral blood^1^. Multiple cell types, including endothelial cells, pericytes, and astrocytes, form the tight structure of BBB. This barrier prevents the entrance of pathogens and toxic substrates into the brain, but also presents a major challenge in drug delivery to the brain^2^. Vascular cells also supply neuronal and glial cells with nutrients and remove waste products, dynamically responding to the changing activity-dependent local energy demands stemming from the brain’s large energetic needs and the lack of local energy storage, through tight cellular interactions and communication between neuronal, astrocyte, and vascular cells forming the neurovascular unit that senses neuronal activity, controls blood flow, and maintains BBB integrity.

BBB breakdown may be an early feature of Alzheimer’s disease (AD), preceding dementia and neurodegeneration, suggesting a critical role of neurovascular unit dysfunction in the progression of AD^3,4^. This leads to the entrance of toxic molecules, pathogens, and cells from peripheral blood to the central nervous system (CNS), triggering inflammatory and immune responses^5,6^. Impaired BBB function is associated with multiple neurodegenerative diseases in addition to AD, including multiple sclerosis (MS) and Parkinson’s disease (PD) in human and mouse model studies^6–8^. In fact, the vascular hypothesis of AD proposes brain vascular damage as the initial event catalyzing BBB dysfunction, precipitating brain dysfunction and cognitive decline^9,10^. However, whether AD risk genes regulate vascular function remains poorly understood systematically.

Brain vascular cells have evaded unbiased characterization due to technical challenges in their isolation, and the cellular complexity of the vascular arbor. The combination of single nucleus RNA sequencing (snRNA-seq) technologies and vessel enrichment protocols has provided an atlas of brain vasculature cell types in human and mouse^11–13^. However, cell sorting and enrichment protocols can introduce technical biases in cell type composition, and result in inaccurate spurious cell compositional inferences, especially if the marker genes used for sorting and enrichment change expression in the context of disease, leading to differences in capture efficiency. Thus, molecular characterization of human cerebrovascular cell types using sorting-free and enrichment-free methods is still needed to understand the cellular basis of the neurovascular unit. Moreover, different brain regions have been shown to possess cellular, morphological and functional differences in vasculature^14^. Understanding this molecular heterogeneity can provide insights into the unique vulnerabilities of different brain regions to disease.

Here, we address these challenges and report a single-cell characterization of the human cerebrovasculature in post-mortem samples from 6 brain regions across 220 AD and 208 age-matched control individuals. We use *in-silico* sorting to capture 22,514 cerebrovascular cells in 11 subtypes, including endothelial, pericytes, smooth muscle cells, perivascular fibroblasts, and ependymal cells. Comparing between brain regions, we find substantial differences in cell type proportion and in gene expression patterns between brain regions, highlighting the regional heterogeneity of the BBB for each cell type. Comparing between AD and non-AD individuals, we find 2,676 cell-type-specific differentially-expressed genes, and we predict their upstream regulators, whose targeting may help restore vasculature function. We also find ligand-receptor-supported coordinated co-expressed gene modules between vasculature, glial, and neuronal cells with diverse roles, including axon growth/degeneration and neurogenesis, suggesting mechanistic mediators of neurovascular unit dysregulation in AD. Integration with AD genetics reveals 125 AD-differential genes directly linked to AD-associated genetic variants (through vasculature-specific eQTLs, Hi-C, and correlation-based evidence), 559 targeted by AD-associated regulators, and 661 targeted by AD-associated ligand-receptor signaling. Lastly, we show that APOE4-genotype associated differences are significantly enriched among AD-associated genes in capillary and venule endothelial cells, and subsets of pericytes and fibroblasts, which underlie the vascular dysregulation in APOE4-associated cognitive decline.

## Results

### Brain vasculature characterization across six brain regions

To characterize human cerebrovascular cells and their transcriptomic differences in AD at single-cell resolution, we profiled and analyzed the transcriptome of 22,514 single nuclei from 725 *post mortem* brain samples of 220 AD and 208 control individuals (contributing 10,272 and 12,242 nuclei respectively, **Supplementary Table 1**) across 6 brain regions, selected using *in silico* sorting using both known marker genes and *de novo* clustering (**Methods**)^12^. We profiled the prefrontal cortex for 409 individuals, and five other regions for a subset of 48 individuals, including mid-temporal cortex, angular gyrus, entorhinal cortex, thalamus and hippocampus, as well as hippocampus for an additional 19 individuals (**Supplementary Table 1**).

We annotated 11 vascular cell types, including three types of endothelial cells (marked by FLT1, CLDN5), two types of pericytes (marked by RGS5, PDGFRB), two types of smooth muscle cells (SMCs) (marked by ACTA2), three types of fibroblasts (marked by COL3A1), and ependymal cells (marked by TTR) (**Fig. 1a-b**), using expression of canonical markers^11,12^ (**Fig. 1c,d**). Consistent with two recent studies^12,13^, we found distinct transcriptomic signatures of arterial, capillary, and venule endothelial cells, as well as both arterial and venule SMCs, suggesting functional specialization for different types of vessels. The inclusion of ependymal cells by our marker-based *in silico* cell sorting highlights the transcriptional commonalities of cerebrospinal fluid (CSF) and BBB barriers, as ependymal cells are not part of cerebrovasculature, but instead form a thin membrane that lines the ventricles of the brain and the central column of the spinal cord where CSF is produced.

**Figure 1.**
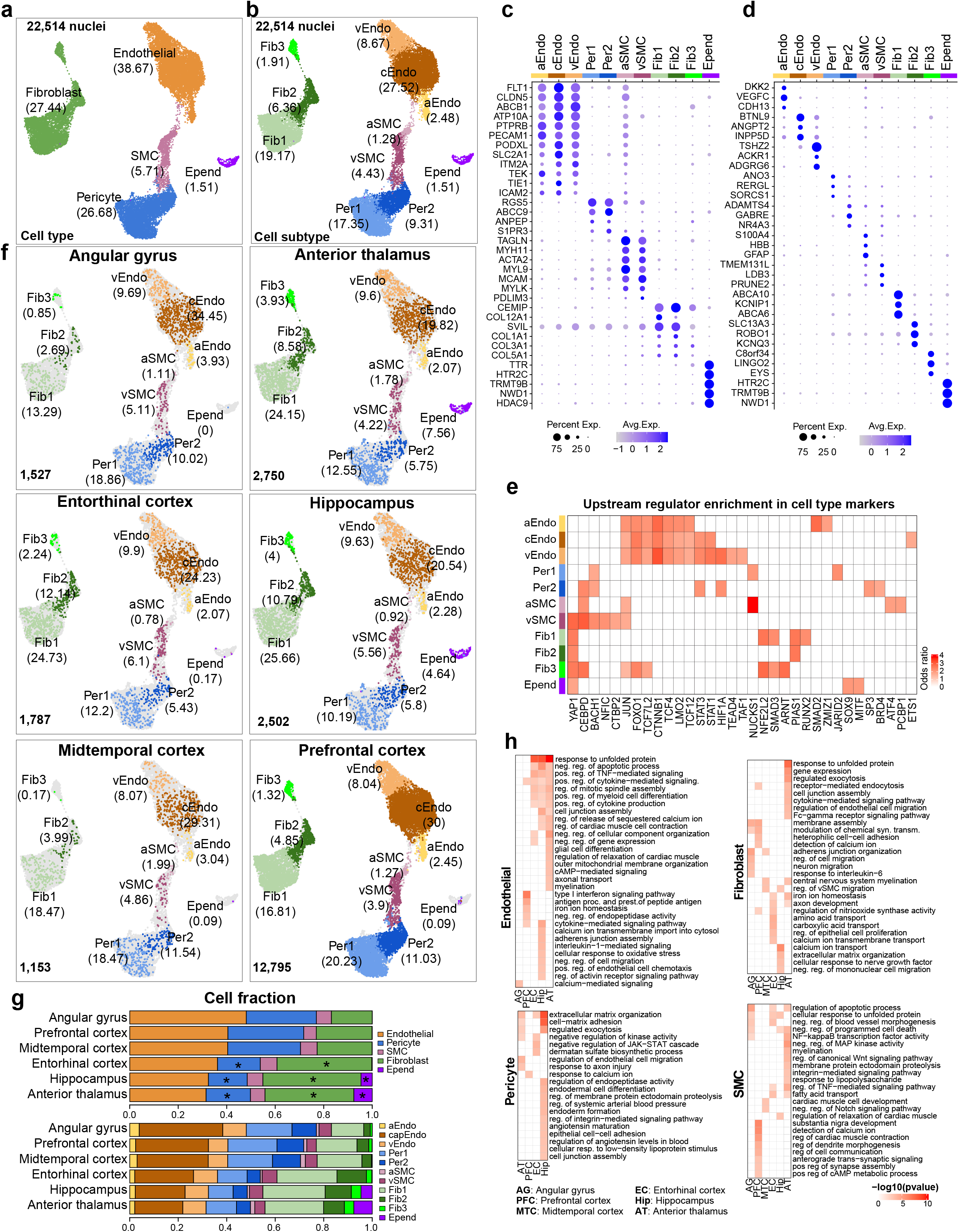
Brain vasculature characterization across six brain regions. **a-b.** UMAP of 22,514 *in silico* sorted brain vascular nuclei from postmortem tissues labeled by cell type (**a**) and cell subtype (**b**), the percentage of cells in each cell is shown. **c-d.** Top markers for vascular cell types (**c**) and cell subtypes (**d**). **e.** Heatmap to show the enrichment of upstream regulators for cell subtype markers. **f.** UMAP of vascular nuclei for each brain region. **g.** Distribution of cell fraction across six brain regions. * represents the significant enriched cell types in specific regions by the Wilcoxon rank test p-value <0.01. **h.** Representative enriched Gene Ontology biological processes of brDEGs for each cell type.

We used transcriptional differences between cell types to highlight key biological functions for each vascular cell type in the brain through pathway-level enrichments of the most highly expressed genes of each cell type (**Extended Data Fig. 1a-b, Supplementary Table 2-3, Methods**). For endothelial cells, the highly expressed genes were most enriched in vascular endothelial growth factor receptor signaling pathway, regulation of cell migration, cytokine response, and cell junction assembly. For pericytes, the highly expressed genes were significantly enriched in pathways including the regulation of angiogenesis, extracellular matrix organization, endothelial cell migration, and cellular sodium ion homeostasis. For SMCs, the highly expressed genes were enriched in smooth muscle contraction, extracellular matrix organization, and regulation of cell junctions. For fibroblasts, the highly expressed genes were enriched in extracellular matrix organization, collagen organization, and regulation of cell migration. In ependymal cells, the highly expressed genes were enriched in cilium assembly, intraciliary transport and ciliary basal body-plasma membrane docking. These results highlight differential functional contributions of brain vascular cells to the maintenance of BBB, regulation of cerebral blood flow, and response to injury.

To gain insights into the gene regulatory programs responsible for establishing cerebrovascular-cell diversity, we next predicted upstream regulators whose activities are associated with the statues of molecular function and cell identity. We performed regulator enrichment analysis for marker genes of each cell type and found that the major cell types tended to share upstream regulators, but still show subtype specificity (**Fig. 1e**). Among the top regulators of cell identity, we found that some regulators show high enrichment in endothelial cells, including CTNNB1, which is associated with maintenance of BBB integrity through endothelial β-Catenin signaling^15^; LMO2, associated with endothelial cell migration in developmental and postnatal angiogenesis^16^; and ETS1, associated with endothelial cell survival in angiogenesis^17^. For pericytes, we found enrichment of BACH1, consistent with its transcriptional regulation of pro-angiogenic activity via modulating the expression of angiopoietin-1^18^. We also found that the AD associated gene YAP1 is a common regulator in vSMC, fibroblast and ependymal cells,consistent with its roles in vSMC phenotypic switch^19^, fibroblast differentiation^20^, and ependymal integrity^21^.

We next evaluated whether vascular cells show differences in abundance across brain regions and phenotypic variables. Fibroblasts were enriched in entorhinal cortex, hippocampus and thalamus (**Fig. 1f-g**), consistent with increased vascular fibrosis and calcification of hippocampus and entorhinal cortex associated with aging^22–25^, and basement membrane and extracellular matrix regional differences^26^, likely stemming from fibroblast-secreted collagen and other proteins (**Fig. 1h**). Ependymal cells were largely captured from the hippocampus and thalamus (**Fig. 1f-g**), consistent with these brain regions’ proximity to CSF ventricles^27^. Hippocampus, thalamus, and entorhinal cortex also showed fewer pericytes and capillary endothelial cells, consistent with their paucity of small vessels, as those are locations where large vessels enter the brain^28^. By contrast, vascular cell fractions did not differ by sex, AD pathology, age, or post-mortem interval (PMI) (Wilcoxon Rank Sum test, p-value < 0.05, **Extended Data Fig. 1c-j**).

We found that vasculature cells showed extensive gene expression differences between brain regions, with many region-specific pathway enrichments, highlighting the regional heterogeneity of the BBB, and importance of single-cell multi-region characterization of the cerebrovasculature (**Fig. 1h**). We found 1,636 differentially-expressed genes between brain regions (brDEGs) (**Supplementary Tables 4**), including: 230 endothelial brDEGs, enriched in molecule transport, regulation of cell migration and junction assembly; 491 pericyte brDEGs, enriched in calcium ion response in prefrontal cortex, and arterial blood pressure, cell junction assembly and response to low-density lipoprotein stimulus in hippocampus; 529 fibroblast brDEGs showing region-specific functional enrichments including extracellular matrix organization, exocytosis, immune response, ion homeostasis, and cell migration regulation; and 491 SMC brDEGs enriched in cell communication in prefrontal cortex, myelination in thalamus, muscle relaxation in hippocampus, and Notch signaling regulation in mid-temporal cortex. Several pathways were commonly enriched across multiple brain regions, including apoptosis regulation, cytokine response, and myeloid cell differentiation (**Extended Data Fig. 1k, Supplementary Tables 5**).

### Cell-type-specific brain vasculature differences in AD

To investigate the vascular gene expression association with AD pathology, we identified and analyzed 2,676 differentially expressed genes (adDEGs) between AD and control individuals across all cell types (306 on average for each cell type), which is significantly more than expected using permutation analysis (t-test p-value=0.007, **Extended Data Fig. 2a**, **Methods**). Of these, 2,142 were unique to only one cell type, 185 (23 genes on average in 8 comparisons) were shared between subtypes of the same cell type (for example, 88 between cEndo and vEndo, and 50 between Per1 and Per2), and 349 (3.8 genes on average in 92 comparisons) were between cell types, highlighting the cell-type-specificity of adDEGs (**Fig. 2a-b, Supplementary Table 6**). This suggests that specialized functions of distinct cell types play unique roles to maintain brain homeostasis and may be dysregulated in AD, while the shared effect across cell types may represent a convergent response to AD. Notably, we found that capillary endothelial cells harbored the highest number of adDEGs, which we recapitulated through a downsampling analysis (**Extended Data Fig. 2b**), suggesting the importance of transcriptional differences in capillary endothelial functions associated with AD pathology (**Fig. 2a**).

**Figure 2.**
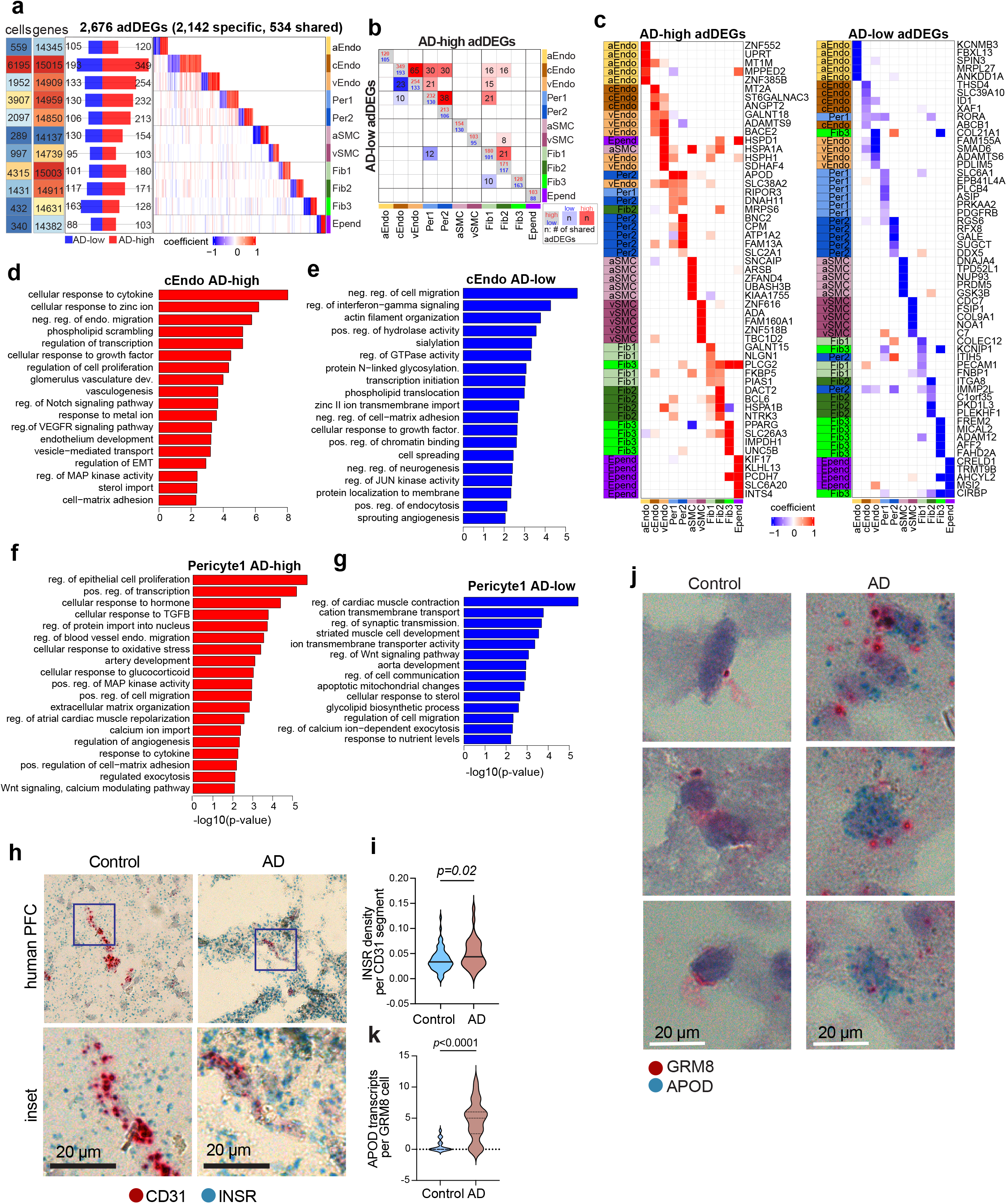
Cell-type-specific brain vasculature differences in AD. **a.** Overview of adDEGs in each cell type. From left to right panels, it shows the numbers of cells and expressed genes, the number of lower/higher adDEGs in AD, and the heatmap with each gene in column and each cell type in row. **b**. The number and significance of adDEGs overlap between cell types in both directions (upper triangle: AD-higher adDEGs; lower triangle: AD-lower adDEGs). **c**. Top5 highly/lowly expressed DEGs in AD in each cell type. The highest effect size for each gene is colored by the cell type. **d-g.** Enriched Gene Ontology biological processes in AD-higher (**d**) and AD-lower (**e**) adDEGs in capillary endothelial cells, AD-higher (**f**) and AD-lower (**g**) adDEGs in Per1. **h-i.** Representative images (**h**) and quantification (**i**) of INSR gene expression in CD31^+^ endothelial cells from control and AD prefrontal cortex tissue. **j-k.** Representative images (**j**) and quantification (**k**) of APOD gene expression in GRM8^+^ pericytes from control and AD prefrontal cortex tissue.

Among the top adDEGs, we observed cell junction and adhesion associated genes including APOD, PECAM1, and COLEC12; transporters including SLC38A2, SLC2A1 and SLC6A1; and sterol-import associated genes including RORA, PRKAA2, and PPARG (**Fig. 2c**), indicating that these fundamental functions of vascular cell types may be dysregulated in AD. As expected, we detected lower expression of PDGFRB in pericytes of AD samples (**Fig. 2c**), alluding to pericyte injury and dysfunction of BBB integrity in AD^4,29^. We also found that ABCB1, encoding P-glycoprotein, was significantly lower in capillary endothelial cells (**Fig. 2c**), consistent with observations that individuals with early AD develop widespread reductions in P-glycoprotein BBB function in multiple brain regions^30,31^.

Gene Ontology enrichment analysis of adDEGs showed that multiple broad functional pathways (e.g. immune response, insulin signaling, and vasculogenesis) were shared across cell types, and multiple specific pathways were more cell type specific (**Fig. 2d-g, Extended Data Fig. 2c, Supplementary Table 7, Methods**). Broad terms included: immune response (cytokine, IL-17 signaling, and inflammatory response) enriched in fibroblast, endothelial cell, and pericyte adDEGs; insulin response enriched in cEndo, pericyte, fibroblast, SMC and ependymal adDEGs, suggesting a potential functional link between altered glucose homeostasis, insulin signaling, and AD pathology across multiple vascular cells^32,33^; vasculogenesis, endothelial cell migration and proliferation, cell-matrix adhesion, and ion transport enriched in cEndo and pericyte adDEGs (**Fig. 2d-f**). Specific terms included: in pericyte adDEGs with lower expression in AD, synaptic transmission (**Fig. 2g**), consistent with pericyte loss of neuronal signal sensing in AD^34^, and cytoskeleton remodeling and contraction (e.g., ATP1A2, ANK2, DMD), consistent with impaired blood flow control and perturbed neurovascular coupling in pericytes in AD^35^; in cEndo, Notch signaling regulation and endothelium development, consistent with the potential role of endothelial Notch signaling and dysfunction of angiogenesis in AD^36,37^; in aSMC with lower expression in AD, cellular response to amyloid-β, consistent with findings showing that amyloid-β deposits contribute to vascular alteration in AD^38^.

Notably, our analysis specifically revealed that insulin signaling genes were AD-differential across multiple cell types (cEndo, pericyte, fibroblast, SMC and ependymal, **Extended Data Fig. 2c**) and in multiple brain regions. Insulin’s cognate receptor *INSR* is widely expressed throughout the mammalian brain, including the hippocampus^39^, cortex^40^, olfactory bulb^41^ and hypothalamus^42^, and research has found aberrant insulin signaling in AD and related dementias^43–45^. To validate our snRNA-based observation that the insulin receptor INSR was highly expressed in endothelial cells from persons with AD, we performed *in situ* hybridization (**Methods**), and found CD31^+^ vascular segments harbored a higher density of INSR1 transcripts in those with AD. We also quantified the distribution of INSR^+^ punctae per CD31^+^ segment, and found a subset of CD31^+^ endothelial cells from AD brains possessing higher INSR transcripts (**Fig. 2h-i, Extended Data Fig. 2d-e**). These results validate our observation that INSR is highly expressed in endothelial cells in AD, and suggest that subsets of endothelial cells may express differential levels of the insulin receptor, potentially rendering cells more or less sensitive to insulin signaling. The degree of heterogeneity of insulin receptor expression in vascular cells, and potential consequences to the neurovascular unit, remain an open question. As sensitivity to insulin signaling is an evolutionarily-conserved mediator of longevity^46^, our results provide further evidence that disruption to insulin is a pathological feature of AD.

We also observed that APOD, encoding secreted glycoprotein Apolipoprotein D, a component of high-density lipoprotein (HDL), is higher in AD samples in mural cells (particularly pericytes, P < 3.9e-12, **Fig. 2c**), in accordance with the previously-reported upregulation of APOD in AD^47^. To validate its higher expression in AD pericytes, we quantified APOD transcript abundance in GRM8-labeled cells (a previously-validated marker for pericytes^12^) using *in situ* hybridization and observed that APOD expression was indeed higher in GRM8^+^ pericytes in AD individuals (**Fig. 2j-k, Extended Data Fig. 2f**) – we did not use PDGFRB as the pericyte marker as it is lowly expressed in AD. Given the function of APOD in response to stress and injury in CNS^48^, the higher expression of APOD in AD pericytes suggests that mural cells are actively responding to microenvironmental changes during AD. The functional consequences of APOD’s pericyte higher expression, and potential impacts on lipid transport within the neurovascular unit, will form the basis of future studies.

Our results show no significant change in vascular cell type proportion between non-AD and AD individuals (**Extended Data Fig. 2g**), and in fact a modest but non-significant high in the median number of capillary endothelial cells and pericytes in AD individuals. These results are consistent with an observed higher proportion of endothelial cells in AD in two recent studies^49,50^ that also did not rely on any cell-type- or vessel-enrichment protocols. However, we found that several our adDEGs that showed lower expression in AD were cell type marker genes, including PDGFRB (pericytes), ABCB1, ATP10A, PTPRB and TEK (capillary endothelial cells), suggesting potential loss of vascular cell type integrity in AD. Such loss of vascular integrity, and lower expression of cell-type-specific markers, could result in a seeming decrease in vascular cell proportions in studies that rely on enrichment protocols.

### Upstream regulators of adDEGs

To gain insights into the transcriptional regulatory mechanisms, we inferred the upstream regulators of adDEGs, including transcription factors, co-factors, and epigenetic enzymes (**Methods, Supplementary Table 8**). We identified 118 upstream regulators of highly expressed adDEGs (60 cell-type-specific, 58 shared by at least two cell types) (**Fig. 3a**) and 81 regulators of lowly expressed adDEGs (41 cell-type-specific, 40 cell-type-shared) (**Fig. 3b**), of which 66 targeted both highly and lowly expressed adDEGs across cell types in AD. Of these 133 regulators, 17 were themselves significant adDEGs corresponding to the differential direction of their targets.

**Figure 3.**
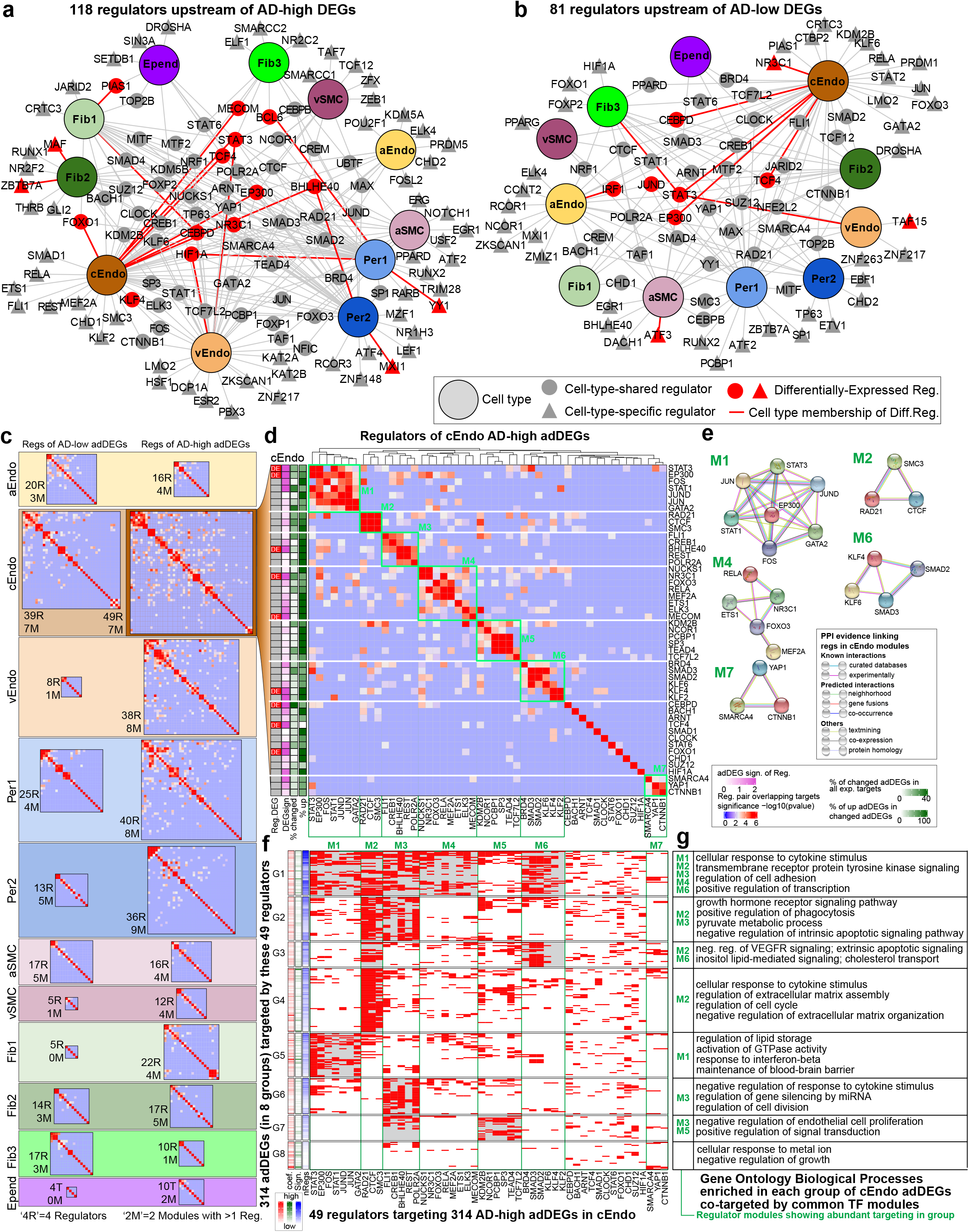
Upstream regulators of adDEGs. **a-b.** Regulator-cell type networks in AD-higher (**a**) and AD-lower (**b**) adDEGs. The large nodes represent cell types. Triangle nodes represent cell type specific regulators. Grey nodes represent regulators with expression. Red nodes represent differentially expressed regulators in at least one relevant cell type. Red edges represent the corresponding differentially expressed regulators in relevant cell types. **c**. Regulator modules of all adDEGs sets. The size of heatmaps reflects the number of regulators. **d**. Regulator modules of AD-higher adDEGs in capillary endothelial cells. Seven modules were labeled by a green rectangle. The first column on the left shows if the regulator is significantly differentially expressed. The second column shows the level of differential significance represented by −log10(p-value). The green columns show the percentage of changed targets for each regulator and the percentage of AD-higher adDGEs in changed targets. **e**. Protein-protein interaction networks from STRING for each regulator module. **f**. Eight groups of adDEGs that are targeted by regulators shown in (**d**). The gray shaded block highlights the regulation of specific regulator modules on the gene group, which is also shown on the left column of (**g**). The first column on the left shows the effect size of adDEGs. The second column shows the level of differential significance represented by −log10(p-value). The third column shows the number of regulators for each target. **g**. Enriched Gene Ontology biological processes of each group of genes in (**f**).

We next grouped these regulators into co-regulatory modules for each cell type (**Fig. 3c, Extended Data Fig. 3**), when regulators showed significant sharing of target genes. Focusing on capillary endothelial cells (cEndo), which showed the largest number of adDEG and upstream regulators, we identified seven regulatory modules (**Fig. 3d**) encompassing 38 regulators, and 11 regulators outside modules. Although our modules were discovered solely based on their shared target genes, most were additionally supported by independent experimental evidence of interactions^51^ (**Fig. 3e**); for example, Module M1 regulators showed a nearly-complete clique of pairwise interactions, each supported by multiple lines of evidence. Moreover, regulators in the same modules were frequently found to have related functions; for examples, Module M1 regulators were shown to have roles in vascular endothelial cell function, growth, and adhesion^52–60^.

Within each module, regulators varied in their number of target genes, their differential expression in AD, and the differential expression of their targets (**Fig. 3d**, left). For example: in Module M1, EP300 previously linked to the potential roles in AD-related processes^61^, was differentially-expressed in AD (1.18-fold higher, P=0.008), and 119 of its targets were adDEGs, of which 74% were AD-higher vs. AD-lower; in Module 4, MECOM previously linked to high-expression in vascular endothelial cells^61–63^ and AD-association^61,62^, showed 1.37-fold AD higher (P=8.8e-6), 53 adDEG targets, with 100% higher expression in AD; in Module 6, KLF4 previously linked to anti-inflammatory properties in endothelial cells^64^ showed 1.21-fold lower expression in AD (P=8.2e-4), 47 (100% AD-higher) adDEG targets, suggesting the activation of inflammatory response in AD. Not all regulators in each module were themselves DEGs, suggesting that some regulators may act through their collaboration with differentially-expressed regulators; for example, STAT3 and EP300 are significant adDEGs in module M1, but JUN, JUND, and GATA2 are not.

We used these regulator modules to partition their target adDEGs into sub-groups, mediated by distinct combinations of regulator modules (**Methods**). For cEndo AD-higher adDEGs, we found groups targeted by a single module (e.g. G4, G5, G6), and other groups targeted by multiple modules (e.g. G1, G2) (**Fig. 3f**), with distinct functional enrichments in common functional categories (**Fig. 3g**). For example, Module 2 (RAD21, CTCF, SMC3) targeted 58% of AD-higher adDEGs (groups G1-G4), consistent with the known role of these regulators on chromatin structure maintenance^65^. Conversely, adDEG genes in Group1 were targeted by regulators from most modules (M1-M4, M6) and were significantly enriched in cytokine response, cell adhesion and transcription regulation. Genes in Group 5, targeted by Module 1 regulators, were enriched in lipid storage, interferon-β response and blood-brain barrier maintenance. Genes in several groups (G3, G7, G8) were enriched in VEGFR signaling negative regulation, repression of endothelial cell proliferation, and apoptotic signaling pathways, suggesting endothelial injury response in AD, consistent with BBB breakdown.

### Dynamics of cell-cell communications in AD

We next sought to understand the mechanistic basis of AD-associated differences in vascular cell communication with glia and neurons in the neurovascular unit, that senses changing neuronal energy demands and adjusts local blood flow, and how this communication is altered in AD. We predicted bidirectional cell-cell communication between vascular and neuronal or glial cells using AD-associated covariation analysis of biologically-enriched gene modules across 409 individuals (**Extended Data Fig. 4a, Methods**). We identified 301 higher interactions in AD, where one or both interacting modules showed higher expression in AD, and conversely 276 lower interactions in AD, where one or both interacting modules showed lower expression in AD, between vascular cell types (capillary and venule endothelial cells, pericytes, and fibroblast subtype1) and other brain cell types (inhibitory/excitatory neurons, microglia, oligodendrocytes, astrocytes and oligodendrocyte precursor cells) (**Fig. 4a-b, Supplementary Table 9**). We found that the communications from Per1 and cEndo to neurons, microglia and astrocytes dominate the higher cell-cell interactions in AD, while interactions from astrocytes and neurons to cEndo and Fib1 are mainly lower in AD.

**Figure 4.**
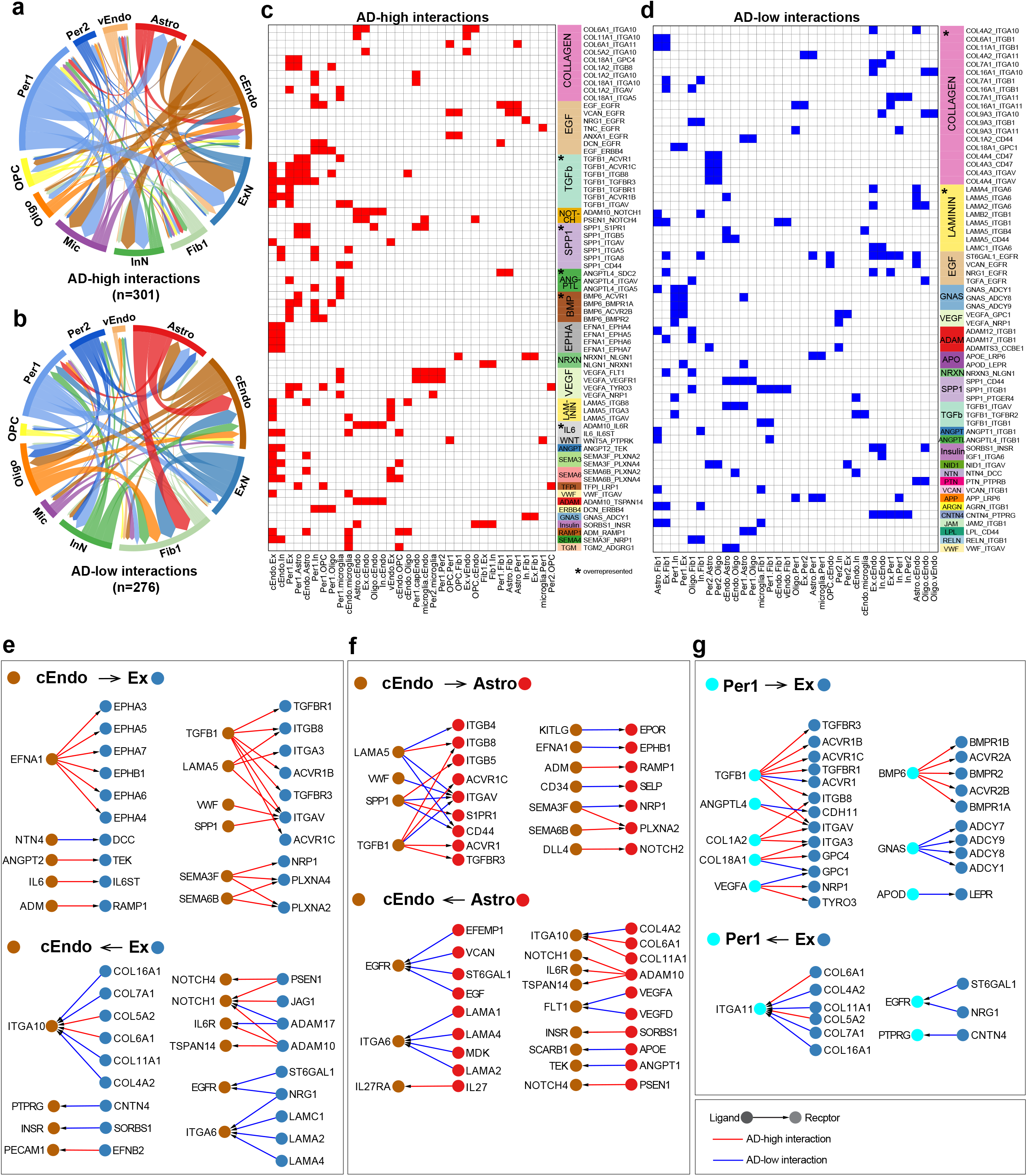
Dynamics of cell-cell communications between vascular cell types and neuron/glial cells in AD. **a-b.** Higher (**a**) and lower (**b**) interactions mediated by ligand-receptor signaling between vascular cell types and neuron/glial/microglial cell types in AD. **c-d.** Ligand-receptor pairs (row) in each pair of cell types (column) for higher (**c**) and lower (**d**) interactions in AD. The signaling pathway is also shown on the right. The star in the signaling pathway column indicates that the signaling pathway is significantly enriched. **e-g.** Top 3 pairs of interacting cell types: cEndo-Ex, cEndo-Astro, and Per1-Ex. Ligand-receptor interactions with direction are shown in the network. The color of nodes represents the cell type. Red edges represent AD-higher interactions, while blue edges are for AD-lower interactions.

To characterize ligand-receptor signaling pathways potentially responsible for differential cell-cell communications in AD, we aggregated individual interactions mediated by the same ligand-receptor pair between vascular cell types and other main cell types, and classified the ligand-receptor pairs into 62 signaling pathways based on KEGG, ligand family, and receptor family annotations (**Methods**). We observed that the signaling pathways mediated by TGF-β, SPP1, BMP, ANGPTL and IL6 are significantly overrepresented for AD-higher interactions, while collagen and laminin, the major ECM proteins to form basement membrane of BBB^66^, are overrepresented for AD-lower interactions (**Fig. 4c-d**), suggesting the potential disruption of BBB structure in AD.

Moreover, we quantified AD-differential interactions in both “forward” and “reverse” communication directions for each pair of cell types and found that interactions between cEndo/Per1 and excitatory neurons/astrocytes show the most interactions (**Extended Data Fig. 4b**), suggesting the important roles of the neurovascular unit in AD pathology. We then built ligand-receptor networks for top three cell-cell pairs (cEndo-Ex, cEndo-Astro, and Per1-Ex) to highlight specific signaling pathways and ligand-receptor pairs (**Fig. 4e-g**). For example, EFNA1, a member of the ephrin family which inhibits axonal growth via EPH signaling^67^, increasingly expressed by cEndo in AD interacts with multiple receptors expressed in excitatory neurons (**Fig. 4e**), suggesting that endothelial cells may mediate axonal growth disruption in AD. TGFB1, which has been shown to exacerbate BBB permeability and regulate pericyte inflammatory response^68,69^, is highly expressed in capillary cell types in AD (cEndo and Per1), and mediates communications with excitatory neurons (**Fig. 4e,g**), suggesting the important roles of TGF-β in reactive oxygen species generation, amyloid-β accumulation and neuronal dysfunction during AD pathogenesis^70^. BMP6 mediates the AD-higher interactions between pericyte1 and excitatory neurons, suggesting its function in pericytes governing impaired neurogenesis in AD^71^. The AD-lower interactions between astrocyte and capillary endothelial cells mediated by EGF signaling pathways suggests inhibition of capillary endothelial proliferation in AD^72^ (**Fig. 4f**).

### AD GWAS loci linked to brain vascular adDEGs

We next sought to gain insight into how AD risk loci may lead to vasculature breakdown at the molecular level, by integration of our adDEGs with AD-associated loci from genome-wide association studies (GWAS)^73–75^ (**Fig. 5a**), to predict their candidate target genes and directionality of effect (higher or lower expression in AD) (**Fig. 5b**), their cell types of action (**Fig. 5c,d**), and the direct (**Fig. 5a-g**) or indirect (**Fig. 5h-j**) mechanisms through which they can lead to vascular gene expression changes.

**Figure 5.**
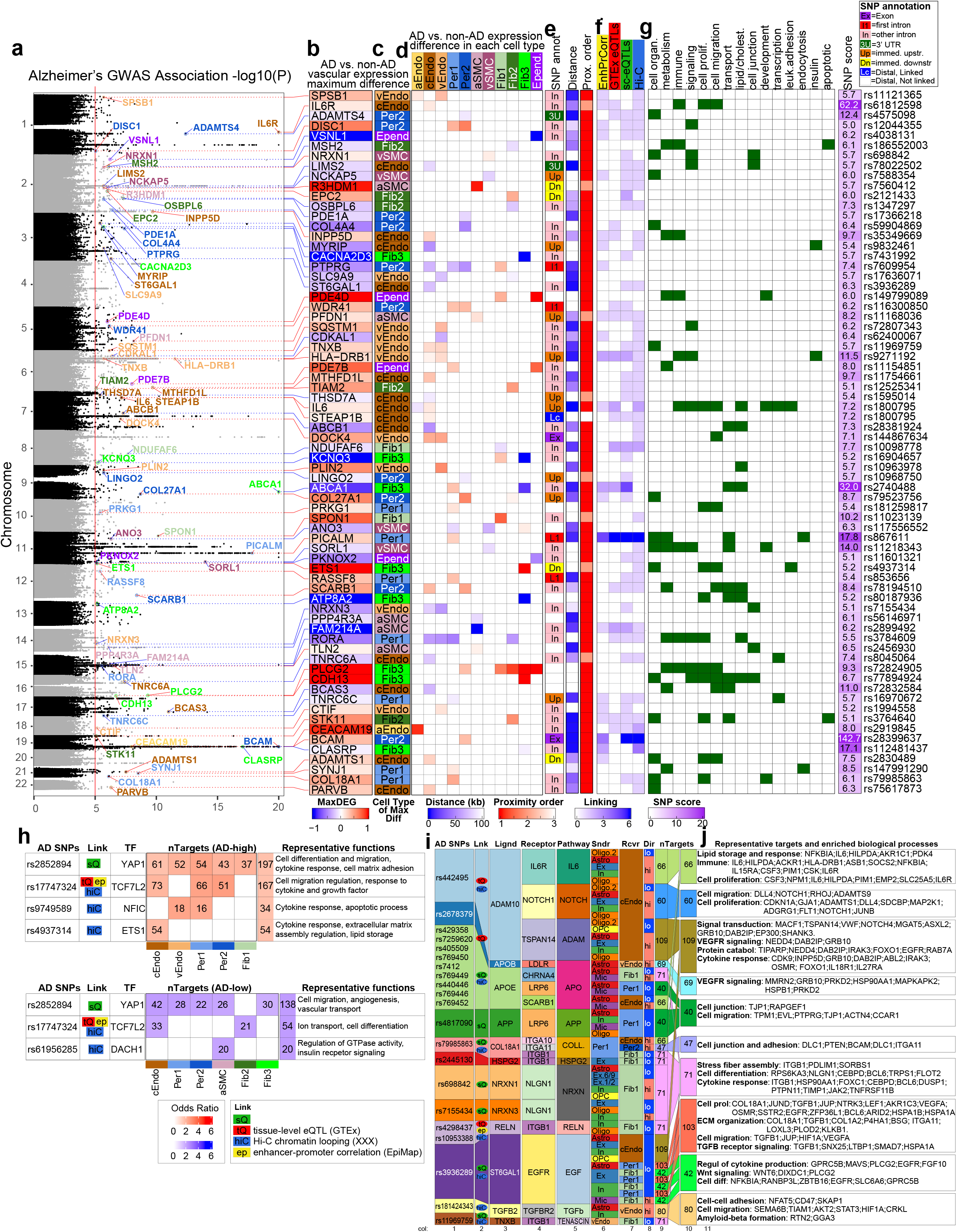
AD GWAS loci linked to brain vascular adDEG. **a-g.** Direct (*cis*) regulated adDEGs by AD-associated variants. Shown are the AD GWAS loci with subthreshold of −log10(p-value) as 5 associated with significant adDEGs (**a-b**), along with the cell-type in which the largest differential expression occurs (**c**), a heatmap to present all differential expression across all cell types (**d**), the genomic annotation for SNPs (**e** left), the distance between variant and adDEG transcription starting site (TSS) (**e** middle), the rank of adDEG among all associated genes of the specific variant (**e** right), a heatmap with four pieces of linking evidence (gene-enhancer correlation from EpiMap, tissue eQTL from GTEx, sc-eQTL, and Hi-C data) (**f**), and a heatmap to highlight the enriched functions of adDEGs (**g**). **h**. Summary of AD-associated transcription factors: AD-variants, linking evidence, the number of targets in adDEGs (up: top; down: bottom) and representative functions of these targets. **i.** Summary of AD-associated ligands: AD-variants, linking evidence, ligand, receptor, signaling pathway, sender cell type, receiver cell type, direction in AD, and number of targeted adDEGs. **j**. The enriched biological processes of targets shown in (**i**).

First, we focused on direct regulation of vascular adDEGs by nearby genetic variants, and found 197 AD-associated variants in 113 loci^76,77^ (P<10^−5^) (**Fig. 5a**) proximal to 125 vascular adDEGs (**Fig. 5b, Supplementary Table 10**) that show cell-type-specific expression alterations (**Fig. 5c,d**). Most adDEGs harbored AD-associated variants within their introns (54.3%), or immediately upstream/down-stream with no intervening gene (7.13%) (**Fig. 5e**), or were linked by: physical chromatin conformation capture (Hi-C) looping^78–81^, correlation-based enhancer-gene links^82^; brain/heart/muscle-specific eQTLs at tissue-level resolution^83^; and vasculature cell-type specific single-cell eQTLs (sc-eQTLs) (Park *et al*, in preparation) (**Fig. 5f**). These AD-associated adDEGs were enriched in similar processes as we reported for cell-type-specific adDEG enrichments, including cholesterol transport, sterol homeostasis, regulation of endothelial cell and vascular associated smooth muscle cell migration, regulation of amyloid precursor protein catabolic process and IL6 mediated signaling (**Fig. 5g**, **Supplementary Table 10**), and we highlight some examples next.

Twenty-one GWAS loci were associated with lipid and cholesterol metabolism adDEG genes, consistent with broad dysregulation of brain cholesterol homeostasis in AD^84^, including several notable examples. First, RORA (**Extended Data Fig. 5b**), a lipid-sensing nuclear receptor, was broadly lowly-expressed (in cEndo, vEndo, pericytes, fibroblasts), was linked (by endothelial Hi-C chromatin loops, muscle eQTLs, SMC sc-eQTLs, pericyte sc-eQTLs) to its own AD-associated intronic variant (rs3784609), and was previously-shown^85^ to regulate pathological retinal angiogenesis by repressing inflammation repressor SOCS3, which is indeed also highly expressed in capillary and venule endothelial cells in AD in our data. Second, ABCA1 (**Extended Data Fig. 5c**), a cholesterol transporter, was linked (by correlation-based enhancer-gene links, muscle and fibroblast eQTLs, sc-eQTLs in endothelial, fibroblast, SMCs) to four AD-associated intronic variants, and was highly expressed in pericytes, which increases pericyte cholesterol efflux to ApoE^86^, and lowly expressed in fibroblasts, which reduces amyloid-β deposition and clearance through ApoE^87,88^. Third, SCARB1 (**Extended Data Fig. 5d**), a cholesterol exchange regulator, was linked (by endothelial Hi-C, endothelial sc-eQTLs, muscle eQTLs) to its own AD-associated intronic variant (rs78194510), was highly expressed in capillary endothelial cells and pericytes in AD, and was shown to mediate HDL signaling in endothelial cells and regulate astrocyte-Aβ and SMC-Aβ interactions^89,90^.

Nine AD GWAS loci were associated with immune response, insulin secretion and neurodegenerative pathogenesis, including several notable examples. First, IL6 and its receptor IL6R were highly expressed in capillary endothelial cells of AD individuals, suggesting immune response of endothelial cells may be a prominent feature of AD blood vessels^91,92^. Second, MYRIP, an insulin secretion regulator, was associated with 100kb upstream variant rs9832461 through Hi-C loop in endothelial cells and enhancer-gene correlation in endothelial cells and brain, and was lowly expressed in capillary endothelial cells of AD (**Extended Data Fig. 5f**), consistent with the expression change in AD^93,94^, and suggesting potential dysregulation of insulin signaling^95^. Third, PFDN1, encoding one subunit of prefoldin complex associated with AD pathogenesis ^96,97^ and showing lower expression in venule SMCs of AD patients, was linked to AD variant rs11168036 through enhancer-gene correlation in brain, heart, and muscle, Hi-C loop in endothelial cells and eQTLs in muscle and endothelial cells (**Extended Data Fig. 5g**), suggesting the dysregulation of protein folding machinery in cerebrovascular cells in AD.

Second, we searched for indirect genetic evidence for adDEGs whose upstream regulators were directly linked to AD-associated variants (**Fig. 5h**). Five of our previously-predicted (**Fig. 3a-b**) upstream transcription factors (YAP1, TCF7L2, NFIC, ETS1, DACH1) were directly linked to AD-associated variants, a 2.9-fold enrichment, given only 33 TFs lie in AD-associated loci (Fisher’s exact test, p-value=0.04). The first four TFs were also predicted to be upstream regulators of cell-type-specific marker genes in our earlier analysis (**Fig. 1e**). These were linked to AD-associated SNPs through diverse lines of evidence: YAP1 through sc-eQTLs in endothelial cells and pericytes; ETS1 through HiC loop in endothelial cells; TCF7L2 through EpiMap promoter-enhancer correlation in endothelial cells and heart, HiC loop in endothelial cells, and eQTL from GTEx in heart; and NFIC and DACH1 through HiC loop in endothelial cells. These AD-associated adDEG regulators targeted 559 vascular adDEGs across five cell types (**Fig. 5h, Supplementary Table 11**), which showed biologically-meaningful enrichments, including: for YAP1, cell migration regulation, angiogenesis and extracellular matrix organization across multiple cell types; for ETS1 cytokine and growth factor stimulus response in capillary endothelial cells, consistent with prior work^17,19–21^; for TCF7L2, a known master regulator in vascularization, SMCs plasticity, glucose homeostasis and insulin production and processing^98–100^ in cEndo, pericytes and fibroblasts.

Third, we searched for indirect genetic evidence for vascular adDEGs downstream of ligand-receptor pairs, whose ligands are directly linked to AD-associated variants, and differentially expressed in AD (adDEGs) in nonvascular cells (neurons, astrocytes, oligos, OPCs, microglia), thus potentially leading to vascular cell dysregulation through ligand-receptor signaling pathways. We used our previously-annotated correlated module pairs (**Fig. 4**, **Extended Data Fig. 4**) that showed coordinated expression differences in AD between vascular and neuronal or glial cell types, and searched for AD-linked adDEG ligands indicative of potential genetic effects. Among our 577 previously-defined module pairs, we found 54 pairs (24 AD-higher and 30 AD-lower) with ligands proximal to AD-associated genetic loci (**Fig. 5i**), implicating 611 vascular adDEGs, which are linked to 12 AD-associated ligands and 13 receptors (in 18 different ligand-receptor pairs). The 12 AD-associated ligands represent a significant enrichment over expectation (odds ratio=2.52, p-value=0.0016, Fisher’s exact test). Of these 12 ligands with proximally linked AD-associated variants, 7 showed sc-eQTL linking evidence, 3 showed tissue-level brain-eQTLs evidence from GTEx, 7 showed Hi-C loop linking evidence, including 6 with multiple lines of evidence (**Fig. 5i**, col. 2). Their downstream genes were enriched in at least 17 different biological functions (**Fig. 5j**, **Supplementary Table 12**). For example, AD-associated rs442495 (P-value=3e-11) in Chr15p13 was linked (through HiC loop and tissue-eQTLs) to ADAM10, a key modulator of dendritic spine formation and AD pathology^101^, an adDEG differentially expressed in multiple cell types (oligodendrocyte, astrocyte, excitatory and inhibitory neurons, and OPCs), that binds three adDEG receptors (IL6R^102^, NOTCH1^103^, TSPAN14^104^), all differentially expressed in capillary endothelial cells, and targeting 66, 60, and 109 receptor-downstream target genes, respectively. The 66 genes downstream of ADAM10-IL6R signaling activates high expressed endothelial genes enriched in lipid storage and response, immune response, and cell proliferation. Interacting with NOTCH1 induces genes involved in cell migration and proliferation. We also observed APOE, the strongest AD genetic associated gene and showing higher expression in microglia and lower expression in astrocyte, interacts with LRP6^105^ to regulate the expression of genes in pericytes significantly enriched for cell junction and cell migration, which is lower in AD.

Taken together, we found 1,010 of 2,676 vascular adDEGs in AD can be associated with AD genetics using *cis, trans-*, or signaling regulatory mechanisms (**Extended Data Fig. 5a, h, Supplementary Table 13**). The expression and transcriptional differences in vascular cell types in the context of AD point out that the effects of genetic risk factors on cerebrovasculature may also contribute to the pathogenesis of AD through the intracellular dysfunction and underlying intercellular communications with neural, glial and microglial cells in the brain parenchyma and immune cells in the peripheral blood system.

### APOE4-associated transcriptional differences and cognitive decline

The apolipoproteinE (APOE) genetic locus is the largest genetic risk factor of late-onset AD with an increased risk for APOE ε4 allele carrier (E4) relative to the common ε3 allele^106^ (E3), capturing more AD heritability than all other known markers combined^107^. APOE ε4 has been previously reported to exacerbate BBB breakdown and pericyte dysregulation^108^, which are thought to contribute to cognitive impairment^109–111^. To elucidate the molecular mechanisms and vascular cell types potentially mediating the effects of APOE ε4 on BBB dysfunction and cognitive decline, we searched for APOE-genotype-associated differentially expressed genes (apoeDEGs) between cells of APOE ε3|ε3 homozygous individuals (E3, N=251) vs. carriers of one or two APOE ε4 alleles (E4, N=101 heterozygous ε3|ε4 and N=7 homozygous ε4|ε4), controlling for other pathological and demographic variables, including AD, age, sex, PMI, Parkinson’s, Lewy Body, VCID, and cognitive decline (see Methods). As the vast majority of ε4 carriers are heterozygous in the population (and in our cohort), we do not focus on homozygous carriers here.

We found 2,482 apoeDEGs (**Fig. 6a, Supplementary Table 14, Methods**), which were mostly evenly distributed across cell types (120 APOE4-higher and 120 APOE4-lower on average) and similar in count to the number of adDEGs. While only a median of 4% of APOE-differential genes were shared with AD-differential genes, reflecting the additional contributors to AD beyond the APOE genotype, the overlap between apoeDEGs and adDEGs was highly significant for a subset of cell types. For capillary endothelial cells (cEndo), 36% of APOE4-higher apoeDEGs were also adDEGs (12-fold enrichment, P=10^−32^, Fisher’s exact test) and 21% of APOE4-lower apoeDEGs were also adDEGs (12-fold, P=10^−20^). Strong apoeDEG-adDEG agreement was also found for Fib1 and Per1 for both APOE4-higher and APOE4-lower genes (10-fold to 14-fold enrichment), and for weaker ones for Per2 and vEndo APOE4-higher genes (6-fold to 8-fold), but no significant overlap was found in other cell types.

**Figure 6.**
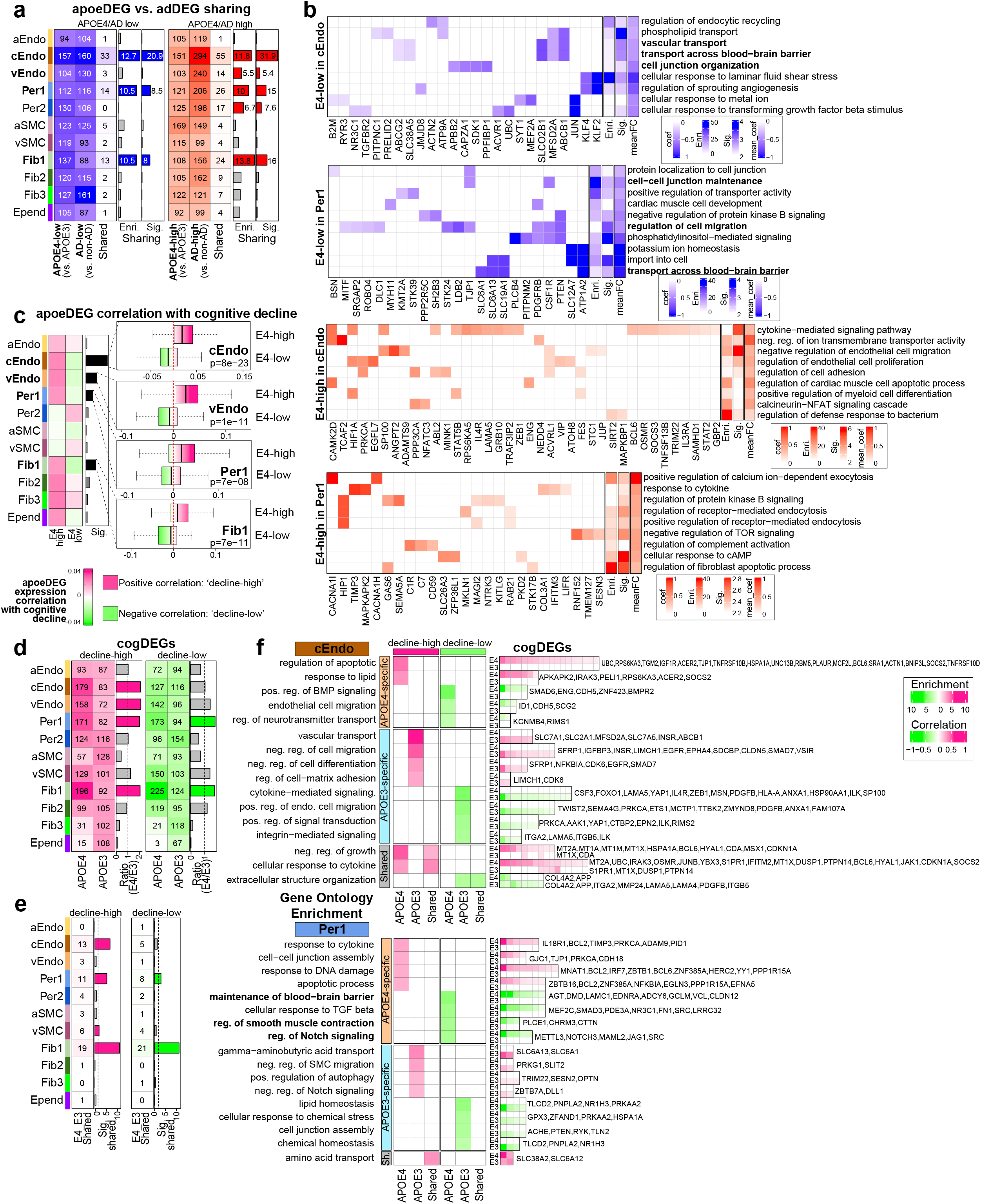
APOE4-associated transcriptional differences and cognitive decline. **a.** The comparison of apoeDEGs and adDEGs for each cell type. The heatmaps show the number of apoeDEGs, adDEGs and overlapping DEGs. The fold enrichment and FDR are shown in the barplot along with heatmap. The significant ones are colored by blue or red. **b**. Enriched Gene Ontology biological processes in E4 lower/higher genes in cEndo and Per1. The color for heatmap of term-gene pairs represents the effect size. The last three columns show enrichment/significance of each term and average effect size of the genes. **c**. The average correlation of apoeDEGs with cognitive decline shown in the heatmap. The bar plot shows the significance of lower correlation of E4 highly expressed genes with cognitive decline.The boxplot (right) highlighted the most significant ones. **d.** The number of decline-higher genes (left) and decline-lower genes (right) in APOE4 and APOE3 individuals. The bar plot shows the cell types with more cogDEGs in APOE4 individuals (>1.5 fold). **e**. The number and significance of overlapping decline-higher/-lower genes between APOE4 and APOE3 individuals. **f**. Enriched biological functions of APOE genotype specific/shared decline-higher/-lower genes in capillary endothelial cells (top) and pericytes (bottom).

To gain more insights on the specific biological functions affected by the APOE ε4 genotype, and thus potential therapeutic hypotheses against AD-associated BBB breakdown, we searched for enriched biological pathways in E4-higher and E4-lower genes in each of these cell types (**Fig. 6b, Supplementary Table 15**). In capillary endothelial cells, E4-lower genes were significantly enriched in transport across blood-brain barrier (e.g. ABCB1, ABCG2, SLCO2B1, SLC48A5), cell junction organization (APBB2, CAPZA1, SDK1, PPFIBP1) and regulation of sprouting angiogenesis (JMJD8, KLF4, KLF2). and in pericytes E4-lower genes were enriched in cell migration regulation, transport across BBB (ATP1A2, SLC6A13, SLC19A1, SLC6A1), and cell-cell junction maintenance (CSF1R, TJP1) (**Fig. 6b**), in agreement with reports of ApoE ε4 individuals showing reduced cerebral blood flow^112^, and cerebrovascular abnormalities^113^. APOE4-higher genes were significantly enriched for cytokine response in both capillary endothelial cells and pericytes, negative regulation of cell migration in capillary endothelial cells, and apoptotic process and positive regulation of endocytosis in pericytes (**Fig. 6b**), consistent with increased risk for neurodegeneration and pathology in APOE ε4 individuals.

We next evaluated the correlation of all apoeDEGs with cognitive decline in all cell types, and found that higher expression of E4-higher genes was primarily associated with cognitive decline across all cell types, while conversely higher expression of E4-lower genes was primarily associated with cognitive resilience (**Fig. 6c, Extended Data Fig. 6a**). This effect was strongest for cEndo (p<10^−22^), vEndo (p<10^−11^), Fib1 (p<10^−11^), and Per1 (p<10^−8^), with substantial and highly-significant differences in cognitive loss correlation between E4-higher vs. E4-lower genes. Our results are consistent with previous findings suggesting capillary pericytes might mediate the effect of APOE4 on cognitive decline^111^, and indicates that capillary endothelial cells and fibroblasts subtype 1 might play equally or even more important roles, based on both the overlap of apoeDEGs and adDEGs, and the highly significant correlation with cognitive decline in both cell types.

We further identified cognitive-decline-correlated differentially-expressed genes (cogDEGs) for each vascular cell type, for both APOE3 and APOE4 individuals (**Fig. 6d, Supplementary Table 16, Methods**), distinguishing “decline-lower” cogDEGs that are negatively-correlated with cognitive decline vs. “decline-higher” genes that are positively-correlated with cognitive decline. APOE4 individuals showed more cogDEGs than APOE3 individuals (>1.5-fold enrichment) for cEndo, Per1, Fib1, and vEndo (**Fig. 6d**), especially for decline-higher genes (2.1-fold), suggesting specific cerebrovascular cell types that might mediate the contribution of the APOE ε4 allele on cognitive decline. Comparing APOE3-specific cogDEGs vs. APOE4-specific cogDEGs, we found that the two sets were largely distinct, with only ~5% of cogDEGs in common between APOE4 and APOE3 (**Fig. 6e**), suggesting distinct transcriptional changes and potential specific mechanisms of cognitive decline between ε3-only vs. ε4 carriers; however this small number of shared cogDEGs was significant in Fib1, Per1, and cEndo.

To further investigate the functions of APOE3 vs. APOE4 cogDEGs, we performed Gene Ontology enrichment analysis for E3-specific, E4-specific, and E3-E4-shared cogDEGs, both decline-higher and decline-lower, in both capillary cell types (cEndo and Per1) (**Fig. 6f, Supplementary Table 17**). For cEndo decline-higher cogDEGs, APOE4-specific enrichments included lipid and cytokine response, apoptotic process and negative regulation of growth, and APOE3 enrichments included vascular transport, negative regulation of cell migration, differentiation and cell matrix adhesion. For cEndo decline-lower cogDEGs, APOE4-specific enrichments included blood vessel development, positive regulation of BMP signaling and neurotransmitter transport regulation, and APOE3 enrichments included positive regulation of endothelial cell migration and extracellular structure organization. For Per1 decline-higher cogDEGs, APOE4-specific enrichments included cytokine response, DNA damage response and apoptosis, and APOE3 enrichments included negative regulation of SMC migration and Notch signaling, and autophagy. For Per1 decline-lower cogDEGs, APOE3 enrichments included lipid and chemical homeostasis and cell junction assembly, whereas APOE4-specific decline-lower cogDEGs were enriched for BBB-related functions (BBB maintenance, Notch signaling, contraction regulation), suggesting that APOE ε4-dependent cognitive decline may be primarily mediated by Per1 pericytes (**Fig. 6f**).

## Discussion

In this study, we profiled and analyzed the transcriptome of 22,514 single nuclei, identified the molecular signatures and upstream regulators of eleven brain vascular cell types, and characterized region-specific expressed genes and pathways in 428 AD and control individuals. We identified 2,676 AD-associated differentially expressed genes (adDEGs) with strong cell-type specificity, including low expression of some canonical cell-type markers and key genes for BBB integrity in AD (e.g. pericyte marker PDGFRB^4,29^), highlighting the specialized functions of vascular cell types in the maintenance of the brain-blood barrier and its dysregulation in disease. These adDEGs were enriched in multiple biological pathways broadly across cell types, including immune response, insulin response, and vasculogenesis, specifically in single cell types, including synaptic transmission in pericytes and Notch signaling in endothelial cells. These results suggest dysregulation of both common functions and cell type-specific pathways, providing potential clues for both global and targeted therapeutic directions aiming at the AD cerebrovasculature.

We also predicted upstream regulators of adDEGs, and grouped them into collaborative regulator modules, potentially driven by primary regulators. Using these modules, we clustered targeted adDEGs into groups specified by higher-order combinations of these modules, each of which was associated with distinct biological functions. Our analysis provides a general framework for understanding how regulators collaboratively control dysfunctional gene programs in disease and may aid in prioritizing for therapeutic targets to restore the function of AD-differential genes, developing iPSC-derived vasculature, and planning perturbation experiments.

We next investigated differential cell-cell communications of neurovascular units in AD. Methodologically, we introduced a new computational framework by combining covariation analysis of gene co-expression modules between cell types across 428 individuals with ligand-receptor pairs and their corresponding signaling pathways. We identified 577 AD-differential cell-cell communications (301 AD-higher and 276 AD-lower) and investigated the responsible signaling pathways. Notably, we found that collagen and laminin, the basic components of the basement membrane of BBB structure, were significantly mediating AD-lower interactions, suggesting the BBB breakdown in AD. Signaling pathways including TGF-β, SPP1, BMP and IL6 were overrepresented in AD-higher interactions, suggesting the potential neuronal dysfunction, Aβ accumulation and immune response of NVU in AD. The dynamics of multi-cellular interactions in AD calls attention to the development of multicellular in vitro systems and provides a specific point of view to therapy in the future.

Moreover, our study yielded insights on interpreting AD genetic variants from genome-wide human genetics studies in cerebrovascular cell types. We innovatively proposed and studied three types of mechanisms to understand how AD variants are associated with vascular differential genes in AD: (1) direct regulation in a “*cis*” way (125 adDEGs), (2) indirect regulation in a “*trans*” way (559 adDEGs through five AD-GWAS-proximal regulators) and (3) indirect regulation through intercellular signaling pathways (611 adDEGs through 12 differentially expressed AD-GWAS-proximal ligands). Altogether, we observed that 1,010 of 2, 676 (37.7%) adDEGs could be associated with AD genetics, suggesting the importance of understanding the BBB dysregulation in AD from the genetic perspective and provides a paradigm to be widely applied in multiple scenarios including other cell types in AD and other diseases.

Finally, our study shed light on the molecular and cellular basis to understand the association of cerebrovasculature with APOE4-associated cognitive decline. We found APOE4 highly expressed genes in cEndo, vEndo, Per1 and Fib1 were significantly and positively correlated with cognitive decline, suggesting that these could be the major cell types associated with APOE4-dependent cognitive impairment. We further observed that cognitive-decline-associated transcriptional differences were APOE genotype specific. For example, the significant enrichment of BBB functions in cognitive-decline-lower genes was specific to APOE4 pericytes, demonstrating that APOE ε4 allele leads to cognitive decline through BBB dysfunction at both cellular and molecular levels. This offers the potential therapeutic targets regarding BBB functions in specific APOE genotypes on cognition impairment. Given the limited number of vascular cells, especially for rare cell types in APOE2 and APOE4, further targeted studies are needed to comprehensively understand the association and causality among APOE genotype, BBB function, cognitive decline and AD.

Overall, our multi-region molecular atlas of differential human cerebrovasculature genes and pathways in AD provides an important foundation for guiding AD therapeutics, especially for early-stage interventions where the BBB is increasingly recognized to play a central role.

## Methods

### Human brain samples from ROSMAP

Human brain tissues in this study were obtained from the Religious Orders Study and Rush Memory and Aging Project (ROSMAP, each approved by an Institutional Review Board (IRB) of Rush University Medical Center) with informed consent, an Anatomic Gift Act for organ donation, and a repository consent to allow the data to be shared^114^. Quantitative clinical and pathologic phenotypes of AD were used to assess disease severity. These included global cognition proximate to death, and a measure of global AD pathology as well as the molecularly specific beta-amyloid and PHFtau tangles. Controls were defined as individuals with little to no AD pathology, whereas cases included a spectrum of AD pathology. Thus, case status was based solely based on AD pathology and other variables were allowed to freely associate, as previously reported^115–119^.

### Nuclei isolation from frozen postmortem brain tissue and single nuclear RNA sequencing

We isolated nuclei from frozen postmortem brain tissue as previously described^120^. Briefly, to avoid transcriptome changes due to protease processing, we homogenize the tissue in a Dounce grinder in the presence of low-concentration detergent to lyse the cell membrane and release intact nuclei. Lysates are filtered through 40 um cell strainers, and we purify nuclei using density-gradient centrifugation to eliminate cell debris. To prevent RNA degradation, we carry out all steps at 4°C and in the presence of RNase inhibitor (Takara). We use the isolated nuclei for the droplet-based 10x scRNA-seq assay, targeting 10k nuclei for each region of each individual and prepare libraries using Chromium Single-Cell 3’ Reagent Kits v3 (10x Genomics, Pleasanton CA) according to the manufacturer’s protocol. We sequenced pooled libraries using the NovaSeq 6000 S2 sequencing kits (100 cycles, Illumina).

### snRNA-seq data preprocessing

We aligned the raw reads to human reference genome version GRCh38 (pre-mRNA) and quantified gene counts using CellRanger software v3.0.1 (10x Genomics, Pleasanton CA)^121^. The generated cell-gene count matrix was processed using the Seurat R package v.4.0.3^122^. We used a threshold of 500 unique molecular identifiers (UMIs) to select cells, and a cut-off value of 50 cells to select genes for further analysis. We filtered out the cells with more than 10% mitochondrial genes. The gene count was normalized by the total counts for each cell, multiplied by 10000, and then log-transformed. We identified the top 2000 highly variable genes for dimension reduction using Seurat default parameters. We used Harmony for batch correction^123^, and DoubletFinder to estimate doublet score with the parameter of 7.5% doublet formation rate^124^. The cells with high doublet scores (0.2 as cutoff) were discarded for further analysis. After generating clusters, the cluster that shows high expression of markers of two or more cell types was also treated as doublets and removed for further analysis.

### In Silico Sorting to enrich vascular cells and cell type annotation

For the full datasets with all cell types, we first annotated the cell type for each cluster based on the canonical markers of major cell types in the brain (including excitatory and inhibitory neuron, astrocyte, oligodendrocyte, OPC, microglia and vascular cell)^120^ and the enrichment of a large set of markers^125^ in highly expressed genes of each cluster. We next calculated the cell type score for each cell, which was represented by the average expression of a group of markers of each cell type^125^. The cells were selected as vascular cells for further integrative analysis only if (1) the clusters that the cells belong to were annotated as vascular cell types; (2) the cells show the specific high score of vascular cell types (highest score is 2-fold higher than the second score). We had previously reported data from control individuals^12^, and here report data from AD individuals.

### Identification of differentially expressed genes

We used the Wilcoxon rank-sum test in Seurat with default parameters to identify highly expressed genes for each cell type compared with the left cells, and differentially expressed genes between brain regions for each cell type (expressed in at least 25% of endothelial cells and logarithm of fold change is higher than 0.25). For the comparison between AD and control, APOE3 and APOE4, cognitive decline, we applied MAST to measure the statistical significance for each gene based on a linear model^126^. The covariates including number of cells, number of expressed genes, age, sex, PMI, race, other dementia related pathology (Lewy body dementia, parkinson’s disease and vascular contributions to cognitive impairment and dementia) were controlled in the model. The genes with FDR <0.05 and coefficient > 0.02 were selected for further analysis. To confirm that these differences are biological and not statistical artifacts, we permuted the annotation of AD pathology for each individual and identified the adDEGs using the same computational pipeline, and found that the number of adDEGs is significantly higher than expected by chance.

### RNA in situ hybridization

For human postmortem samples, fresh frozen human PFC samples (BA region 9) were embedded in Tissue-Tek OCT compound (Sakura, #25608-930), cut at 10 μm using a cryostat (Leica, CM3050S) and collected on Superfrost Plus slides (Fischer Scientific, #12-550-15). Tissue sections were stored at −80°C until further processing. RNAscope chromogenic 2.5 HD duplex reagent kit (Advanced Cell Diagnostic, #322430) was used to perform RNA in situ hybridization according to the manufacturer’s instructions with the following modifications: tissue was fixed in 4% paraformaldehyde for 30 minutes; 30 minutes were allowed for C2 probe hybridization; overnight at room temperature was used for C1 probe hybridization; and xylene was not used prior to mounting. Probes used in this study include CD31 (Advanced Cell Diagnostic, #548451-C2, red) and INSR (Advanced Cell Diagnostic, #406411, green). Images were acquired using the brightfield settings of a Zeiss LSM 900 microscope.

### RNA in situ hybridization analysis

Images were imported into QuPath (version 0.2.0-m8). Vessel segments were identified based on CD31+ punctae, which formed visually distinct vessel-like segments. To quantify INSR expression within CD31+ cells, we manually counted the number of INSR+ punctae within each vascular segment. To calculate the density of INSR within each vascular segment, we quantified the ratio of INSR punctae within each vascular segment based on square micrometer surface area of each segment, then determined the frequency distribution for each density quantification between AD and control patients.

### Prediction of regulators

We predicted the upstream regulators of cell type markers and adDEGs using Enrichr in R based on three libraries including TRANSFAC and JASPAR, ChEA, and ENCODE TF ChIP-seq data^127–129^. We used adjusted p-value <0.05 as a cut-off to select the significant regulators. We kept the regulators with detected expression in the relevant cell types for further analysis. For the predicted upstream regulators of adDEGs, we tested the significance of shared targets between each pair of them in each geneset, and grouped these regulators into co-regulatory modules based on the hierarchical clustering of significance (-log10 of p-value by Fisher’s exact test). We evaluated the regulators within one module by searching the protein-protein interaction network database STRING 51 and downloaded the generated network as supporting information in this study. We then generated a matrix of zero-one to represent the regulatory relation between regulators and targets, performed hierarchical clustering to separate targets into distinct clusters and calculated the percentage of ones in the block (formed by gene cluster and regulator module) to determine if the target cluster was regulated by a regulator module (30% as a cutoff). One target group could be regulated by zero, one or multiple regulator modules, and vice versa.

### Gene Ontology Enrichment Analysis

We used Enrichr in R to perform enrichment analysis for Gene Ontology biological process (adjusted p-value < 0.05 as a cut-off)^130,131^. The selected terms were manually combined according to the general functions including immune response, cell proliferation, cell migration, etc.

### Prediction of dynamic cell-cell communications in AD

For each cell type (including vascular and non-vascular cell types in our brain datasets), we first clustered adDEGs into co-expression gene modules across all individuals, aggregating single-cell expression for each cell type within each individual. Then, for each pair of cell types, we generated the Pearson correlation coefficient matrix across all module pairs for that cell type pair, and estimated the significance using cor.test function in R. We used a value of adjusted p-value 0.05 as cut-off to select the significant correlated modules between two cell types. For each gene module, we performed Gene Ontology enrichment analysis to measure the importance of the module at the pathway level and removed the gene modules without enriched biological processes and signaling pathways. We combined four ligand-receptor databases (CellChatDB^132^, CellPhoneDB^133^, CellTalkDB^134^, and SingleCellSingalR^135^) to annotate the correlated modules as evidence for interacting cell-cell pairs mediated by ligand-receptor signaling pathways. We manually curated the selected ligand-receptor pairs in this study to remove the mislabeled or intracellular protein-protein interactions through literature searching and filtering. The final results include the interacting cell types, ligand, ligand-involved functions, receptor, receptor-involved functions, potential targets in signal receiver cell type, and direction of cell-cell communication in AD.

### adDEGs association with AD genetics

We firstly downloaded AD-GWAS associated genes based on GWAS catalog annotation^77^ and summary statistics from Jansen et al^73^ using a subthreshold of −log10(p-value)>5 to match genes from GWAS catalog annotation. The adDEGs in AD GWAS catalog were then mapped to AD variants based on the annotation in catalog. The variant with the lowest p-value for each gene was used to label position in the Manhattan plot. We used four types of linking evidence to evaluate the association between AD-variant and candidate genes: (a) physical chromatin conformation capture (Hi-C) looping^78–81^, (b) correlation-based enhancer-gene links^82^; (c) brain/heart/muscle-specific eQTLs at tissue-level resolution^83^; and (d) vasculature cell-type specific single-cell eQTLs (sc-eQTLs) (in preparation). The association with at least two pieces of linking evidence were selected for visualization. The distance between variant and gene TSS and its rank among variant-associated genes were also calculated and visualized.

We also evaluated the association between AD-variant and predicted regulators of adDEGs using the four types of linking evidence mentioned above. We calculated the number of targets for each AD-associated regulator for each cell type and plotted the heatmap to show if the regulation is cell-type-specific or cell-type-shared. Similarly, we evaluated the association between AD-variant and ligands that meditates AD-related cell-cell communications using the four types of linking evidence and performed functional enrichment analysis for each target set to measure the functions of those targets as the downstream of AD-associated ligand in vascular cell types.

## Acknowledgements

We thank the study participants and staff of the Rush Alzheimer’s Disease Center. This work was supported in part by NIH grants AG054012, AG058002, AG062377, NS110453, NS115064, AG062335, NS127187 (M.K., L.-H.T.); AG067151, MH109978, MH119509, HG008155, DA053631 (M.K.); P30AG10161, P30AG72975, R01AG15819, R01AG17917, U01AG46152, U01AG61356, and R01AG57473 (D.A.B.); and the Cure Alzheimer’s Foundation CIRCUITS consortium (M.K., L.-H.T); The JPB Foundation (L.-H.T.); Robert A. and Renee Belfer (L.-H.T.); N.S. was supported by Takeda Fellowship from Takeda Pharmaceutical Company. We thank Carles A. Boix, Lei Hou and Patricia Purcell for scientific suggestions.

## Code Availability

Code used in this study is available upon reasonable request from the corresponding authors.

## Data Availability

Count matrices for all cells analyzed in this study are uploaded with this submission as Supplementary Data and at http://compbio.mit.edu/scADbbb/. Interactive website is linked from http://compbio.mit.edu/scADbbb/. ROSMAP data can be requested at https://www.radc.rush.edu.

## Author Contributions

N.S., M.K., and L.-H.T. conceived and designed the study; M.K. and L.-H.T. supervised the study; N.S. developed the computational framework and conducted data analysis with assistance from Y.P..; H.M., K.G., X.J. and A.P.N performed snRNA-seq profiling. L.A. and M.H.M. performed *in situ* hybridization and quantification with help from A.B.; D.A.B. provided *post mortem* samples and scientific input; and N.S. and M.K. wrote the paper with comments from all authors.

## Extended Data Figures

**Extended Data Figure 1.**
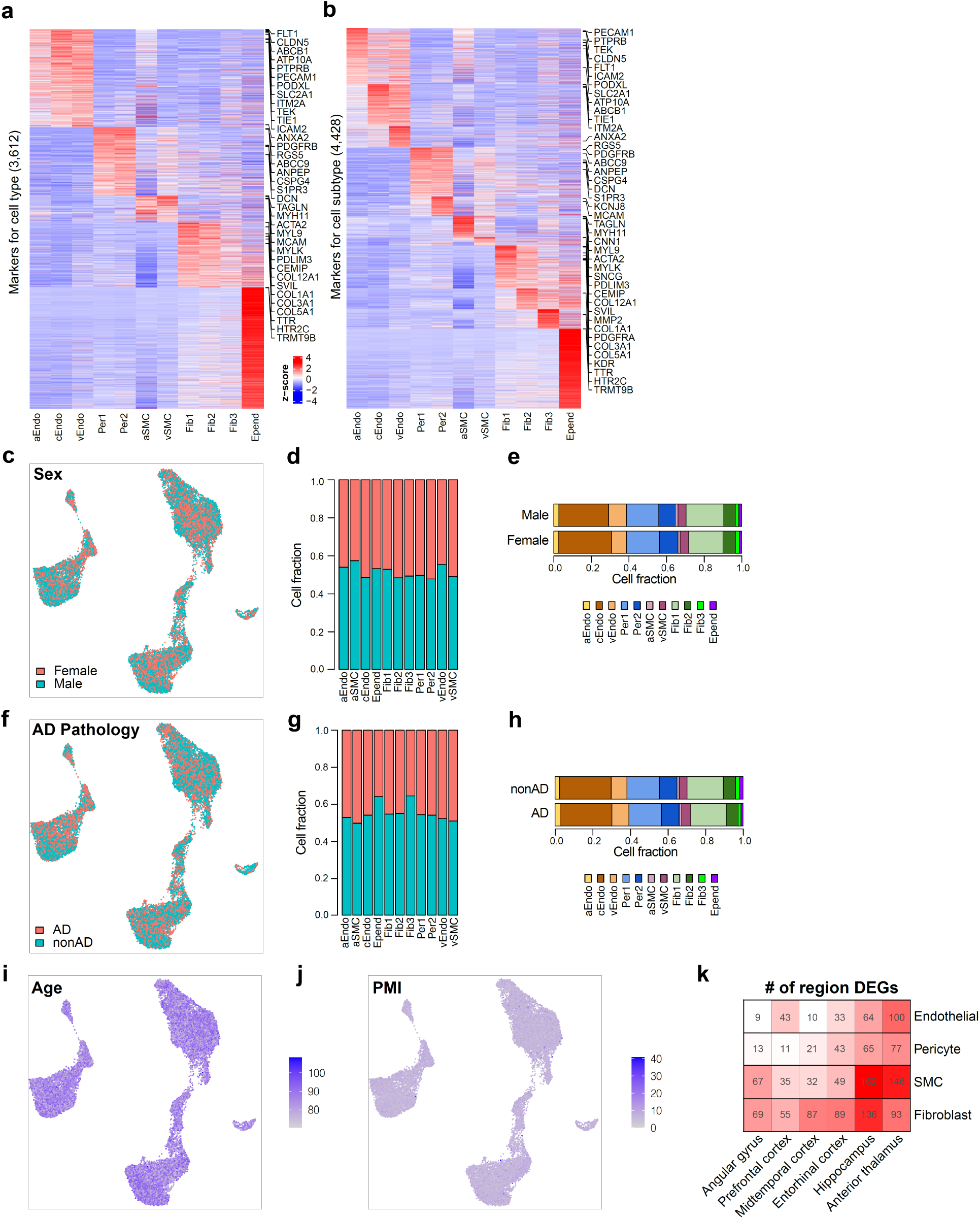
Brain Vasculature Characterization across Six Brain Regions. **a-b.** Markers for vascular cell types (**a**) and cell subtypes (**b**). **c-e.** Cerebrovascular cell distribution by sex. UMAP of brain vascular nuclei labeled by sex (**c**), cell fraction across sex for each cell type (**d**), and cell fraction across cell types for male and female individuals (**e**). **f-h.** Cerebrovascular cell distribution by AD diagnosis. UMAP of brain vascular nuclei labeled by AD diagnosis (**f**), cell fraction across AD diagnosis for each cell type (**g**), and cell fraction across cell types for AD and control individuals (**h**). **i-j.** UMAP of vascular nuclei with age (**i**) and PMI (**j**). **k.** Heatmap to show the number of highly expressed brDEGs of each region for each cell type.

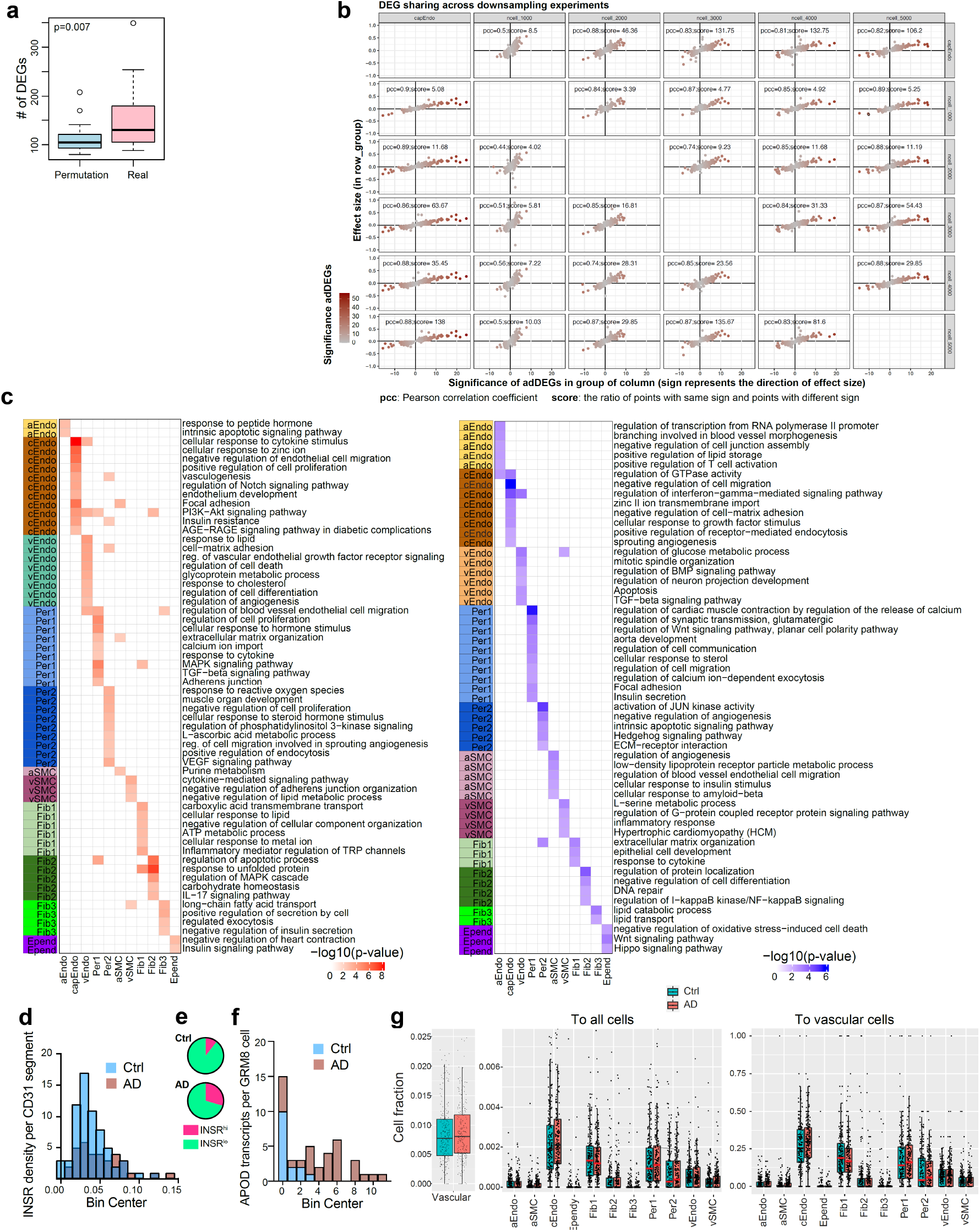

**Extended Data Figure 3.**
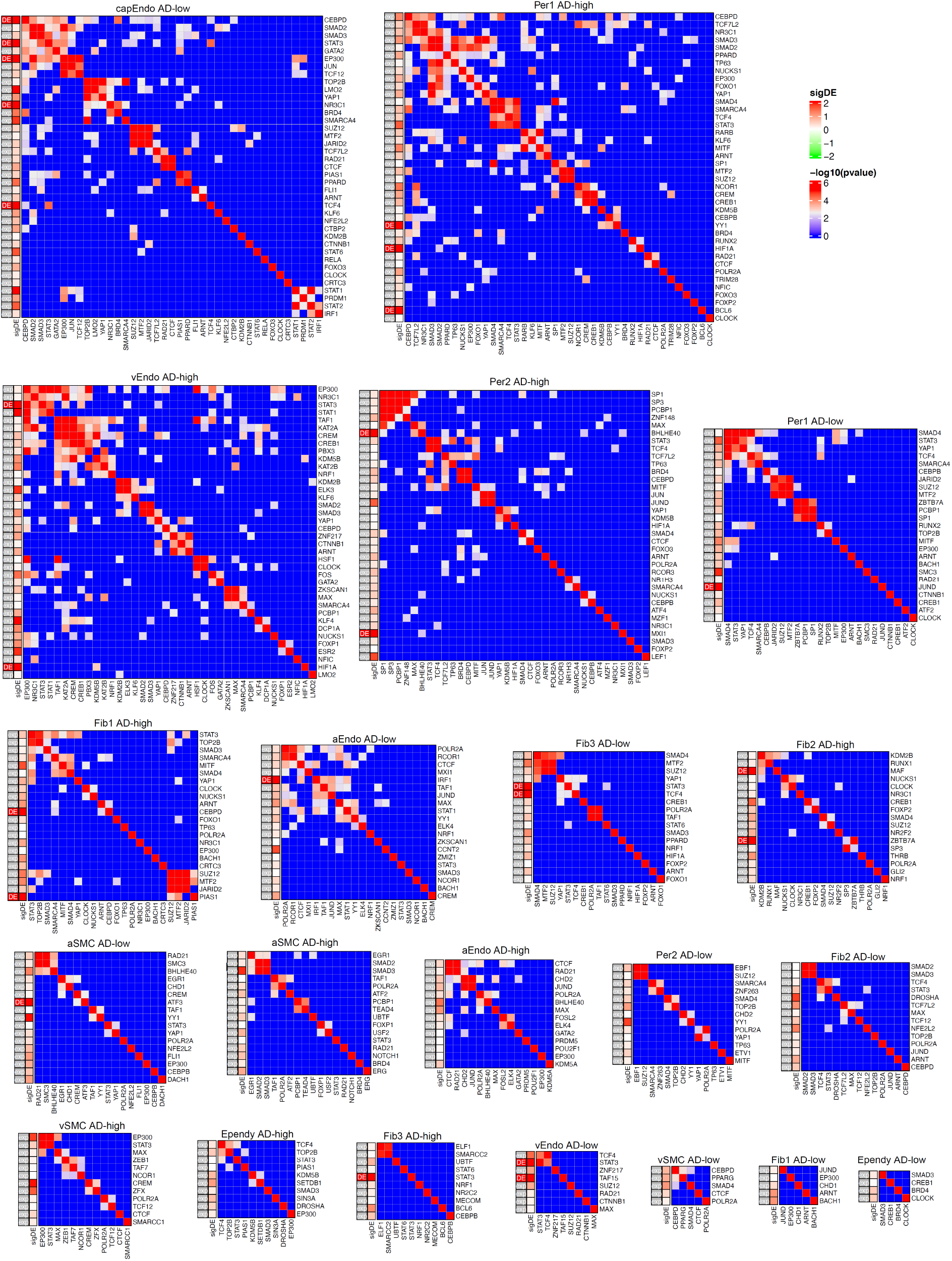
Upstream regulators of adDEGs. Regulator modules of adDEGs in 11 cell types. For each heatmap, the first column on the left shows if the regulator is significantly differentially expressed. The second column shows the level of differential significance represented by −log10(p-value).

**Extended Data Figure 4.**
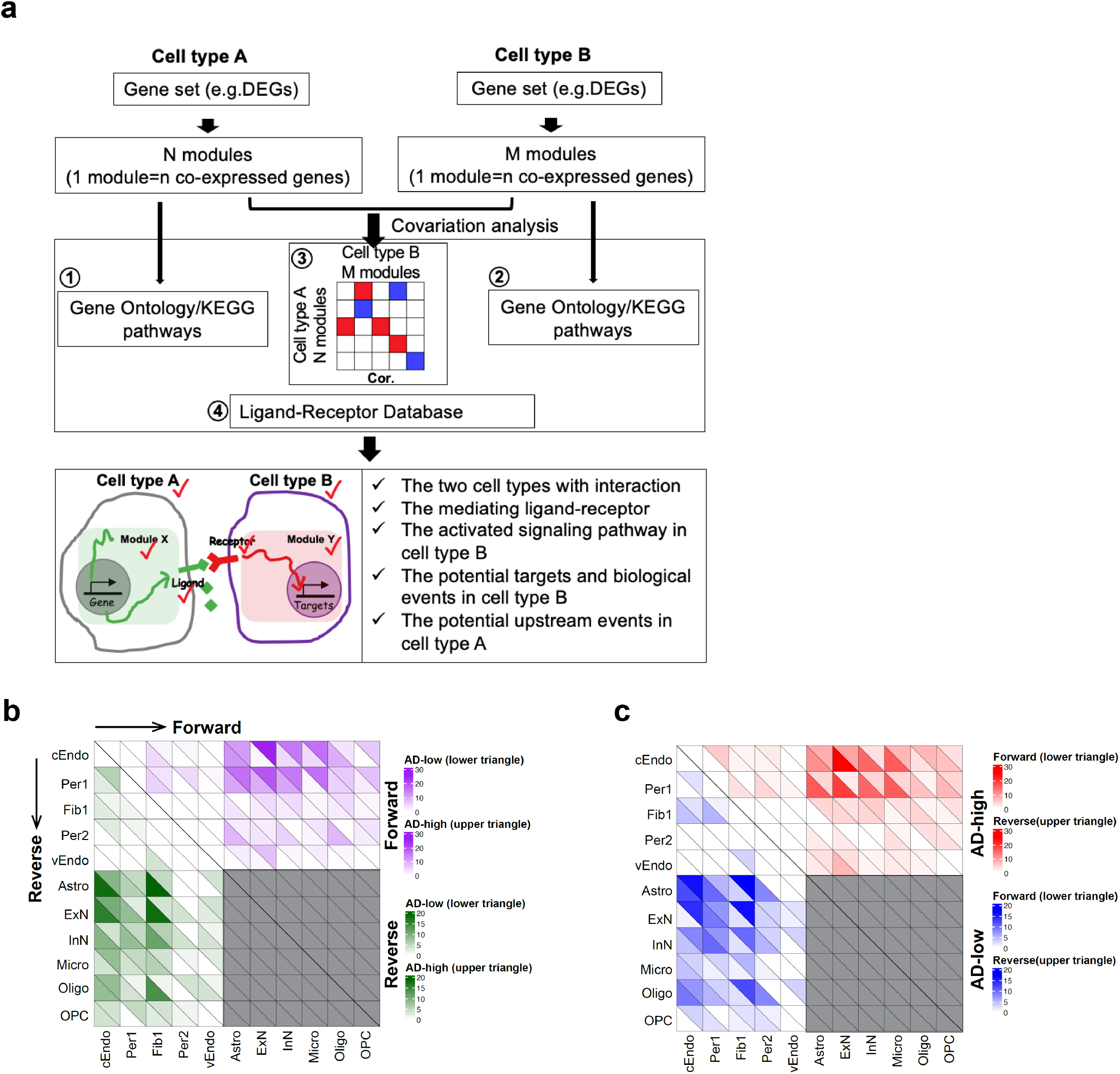
Dynamics of cell-cell communications between vascular cell types and neuron/glial cells in AD. **a.** Computational framework to infer cell-cell communications. For each cell type, a set of genes were clustered into a number of co-expressed modules. The pairwise Pearson correlation coefficient was calculated between modules from each pair of cell types. The significant correlated modules, functional enrichment and ligand-receptor pairs were integrated into the prediction of cell-cell communication. The output includes the interacting cell types, ligand, ligand-involved functions, receptor, receptor-involved functions, potential targets in signal receiver cell type, and direction of cell-cell communication in AD. **b.** The numbers of AD-lower (lower triangle of each square) and AD-higher (upper triangle of each square) in forward interactions from row to column (for example, the first row means the interaction from cEndo to other cell types) (upper triangle of the heatmap) and reverse interactions from column to row (for example, the first column means the interaction from other cell types to cEndo) (lower triangle of the heatmap). **c.** The numbers of forward (lower triangle of each square) and reverse (upper triangle of each square) in AD-higher interactions (upper triangle of the heatmap) and AD-lower interactions (lower triangle of the heatmap).

**Extended Data Figure 5.**
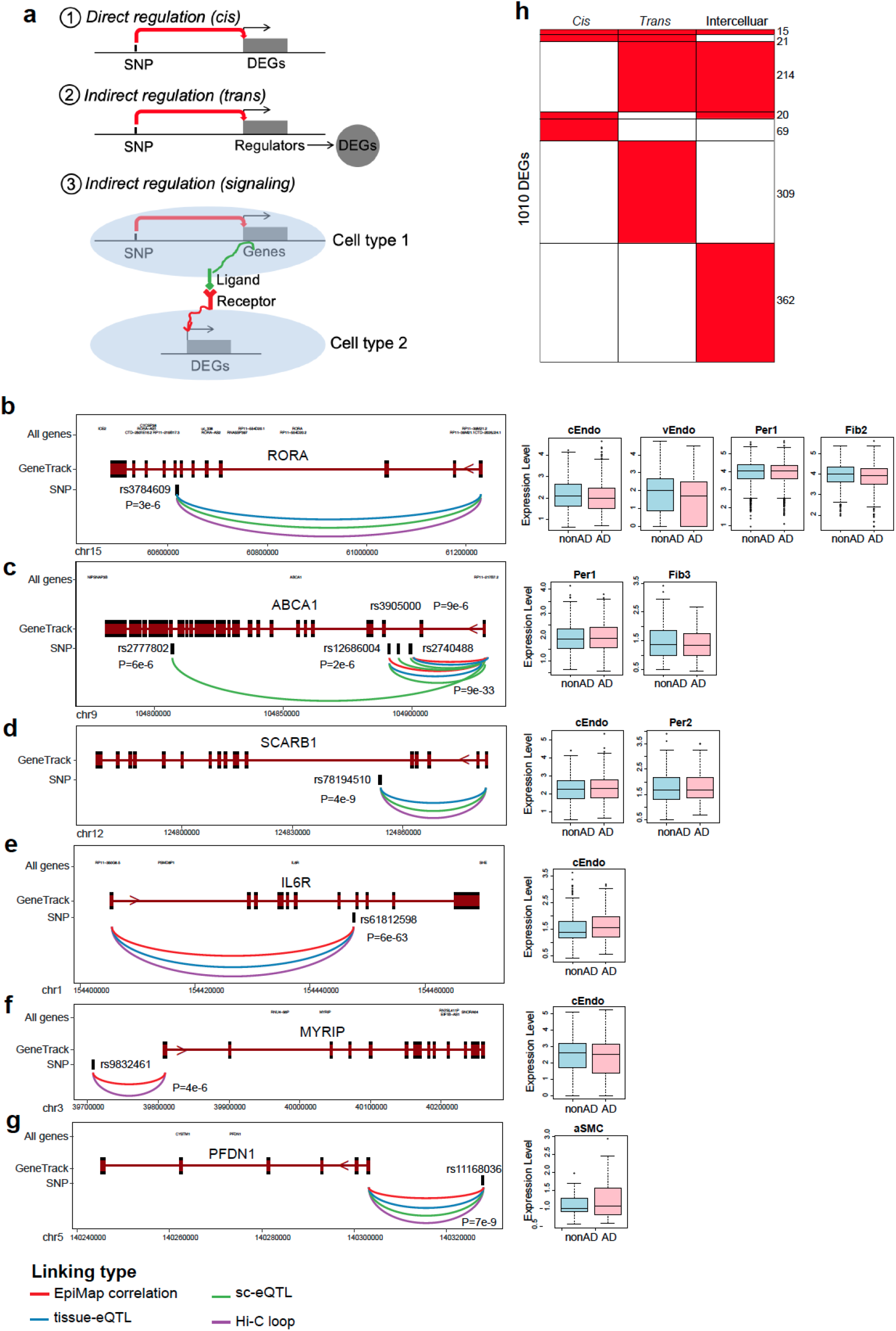
Differentially expressed vascular genes association with AD genetics. **a.** Proposed three types of regulatory mechanisms to interpret the association between adDEGs and AD genetic variants: (1) Directly (*cis*) regulate adDEGs; (2) Indirect (*trans*) regulates adDEGs; (3) Indirect (*ligand-receptor signaling*) regulates adDEGs. **b-g**. Examples of adDEGs directly regulated by AD-associated variants through linking (eQTLs, Hi-C, promoter-enhancer correlation) along with the expression changes in vascular cell types. **h**. The number of targets regulated by the three regulatory mechanisms.

**Extended Data Figure 6.**
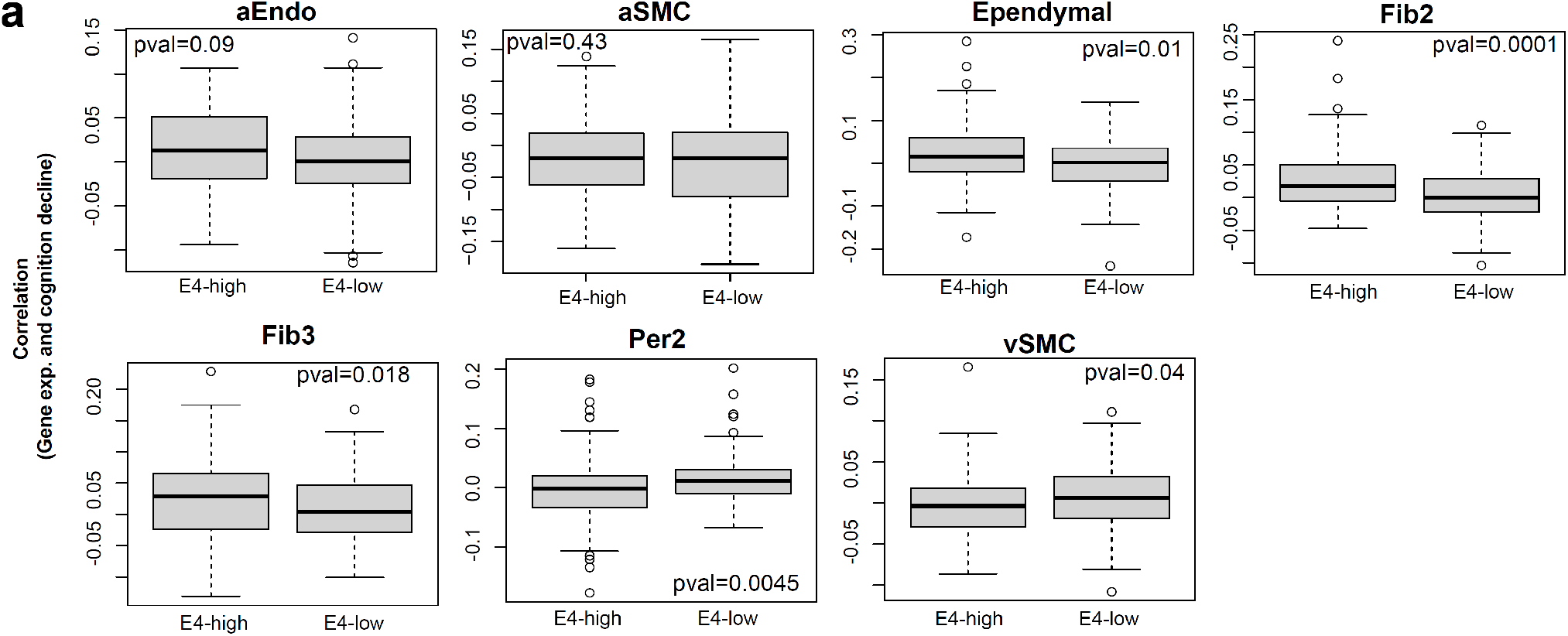
APOE genotype, BBB dysfunction and cognitive decline. **a**. The comparison of apoeDEGs correlation with cognitive decline in each cell type.

## Supplementary Tables

**Supplementary Table 1.** Metadata for ROSMAP samples

**Supplementary Table 2.** Marker genes for cell type and subtypes

**Supplementary Table 3.** Functional enrichment for cell type markers

**Supplementary Table 4.** brain region brDEGs

**Supplementary Table 5.** Functional enrichment for brDEGs

**Supplementary Table 6.** AD adDEGs

**Supplementary Table 7.** Functional enrichment for AD adDEGs

**Supplementary Table 8.** Predicted regulators and their targets

**Supplementary Table 9.** Predicted cell-cell interactions

**Supplementary Table 10.** 125 GWAS genes and their variants, linking evidence, and functional enrichment

**Supplementary Table 11.** GWAS TFs, targets and functions

**Supplementary Table 12.** GWAS ligands, CCI and their targets

**Supplementary Table 13.** Summary of AD GWAS-associated vascular adDEGs

**Supplementary Table 14.** apoeDEGs between APOE3 and APOE4

**Supplementary Table 15.** Functional enrichment for apoeDEGs between APOE3 and APOE4

**Supplementary Table 16.** APOE genotype dependent cognitive-decline-associated genes

**Supplementary Table 17.** Functional enrichment for APOE genotype dependent cognitive-decline-associated genes

